# Guidance of cellular nematics into shape-programmable living surfaces

**DOI:** 10.1101/2025.06.27.660992

**Authors:** Pau Guillamat, Waleed Mirza, Pradeep K. Bal, Manuel Gómez-González, Pere Roca-Cusachs, Marino Arroyo, Xavier Trepat

**Author notes:** These authors contributed equally to this work (Waleed Mirza, Pradeep K. Bal).

## Abstract

Engineering living materials capable of autonomously morphing into predetermined shapes holds great potential for applications ranging from synthetic tissue morphogenesis to soft robotics. In this regard, harnessing the inherent ability of cellular tissues to self-organize and generate robust force fields offers a promising strategy for creating self-shaping living surfaces. However, achieving precise control over tissue mechanics to direct specific morphogenetic outcomes remains a challenge. Here, we introduce a strategy to program tissue shape transformations through the nematic organization of cellular forces. By precisely controlling nematic order and topological defects, we generate millimeter-scale tissues programmed with specific multicellular tension fields. Using a theoretical framework that integrates contractile nematics with thin sheet mechanics, we explore the role of nematically guided tensions in shape morphogenesis. Experimentally, upon tissue detachment, nematically guided tension fields drive out-of-plane deformations, generating reproducible 3D shapes. By integrating tissue contractility and nematic patterning, our approach offers a robust framework for the rational design of shape-programmable living surfaces.

## Introduction

Living surfaces, from the cellular cortex to multicellular sheets, exhibit a remarkable capacity for self-organization and shape change, driven by a finely tuned interplay between molecular signaling and mechanical forces (*1*). At the tissue level, this self-organization manifests as robust supracellular force patterns that sculpt living materials into functional shapes (*2*). Replicating these phenomena in synthetic systems deepens our understanding of tissue morphogenesis and opens new avenues for applications in regenerative medicine, synthetic biology, and soft robotics. Despite significant advances in bottom-up tissue engineering (*3*), our capabilities to precisely program tissue shape transitions are still quite limited (*4–7*). A key challenge lies in directing large-scale force patterns across tissues in a controlled and predictable manner.

A promising approach is to exploit the inherent organizational potential of multicellular systems. Tissues composed of elongated cells often display long-range orientational order, known as nematic order (*8*), analogous to the alignment of rod-like molecules in nematic liquid crystals (*9*). In these systems, hereafter referred to as cellular nematics, the nematic order governs a range of cellular behaviors, from single-cell orientation and migration to multicellular force distributions (*8*). Crucially, local disruption in nematic order gives rise to topological defects, which act as localized sources of force generation (*8*). The configuration of the orientation field around a defect determines its topological charge, a quantity that measures the total angular rotation of the orientation vector along a 2π circuit around the defect (*9*). The topological charge is pivotal in shaping the force fields generated by multicellular topological defects, which serve as mechanical centers for specific biologically relevant functionalities. For example, defects with positive half-integer 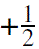 charge can generate localized compressive stresses, which have been linked to processes such as cell extrusion in epithelial layers (*10*) or multilayering in bacterial systems (*11*). Similarly, positive integer defects with charge +1 induce localized compressive stresses that drive differentiation in regions of maximum stress or promote the formation of cellular protrusions (*12*). In addition to compressive stresses, both 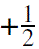 and 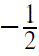 topological defects can concentrate tensile stresses, leading to local drops in cell density (*13, 14*) or even fractures in epithelial layers (*15*). In hydra, topological defects in the supracellular actin network appear to guide morphological traits (*16, 17*), highlighting the importance of topological defects in tissue morphogenesis. In this context, the reshaping capacity of active nematic sheets has been a subject of recent theoretical studies (*18–23*).

Despite these advances, controlled exploitation of cellular nematics to drive predictable tissue morphogenesis remains in its infancy (*12, 24, 25*). A key obstacle has been the lack of integrated strategies to simultaneously control and mechanically probe cellular nematics, which has so far hindered the establishment of a robust link between topological defect forces and shape transformations. Here, we present an experimental and theoretical framework to program 3D tissue shapes through the patterning of 2D nematic force fields. Our approach guides nematic organization while simultaneously enabling the measurement of the forces emerging from collective multicellular interactions. Specifically, we employ anisotropic micropatterns of adhesive proteins to direct cell spreading and migration, forming cellular nematic layers with pre-designed orientational arrangements. By growing fibroblast layers with programmed orientation on soft substrates, we are able to measure the nematically directed supracellular traction and tension fields. Simulations of contractile nematic thin elastic sheets predict how nematic order and topological defects guide tissue shape transformations. Informed by these theoretical results, we gradually detached orientation-programmed cellular nematics from their substrates, reproducibly generating specific 3D tissue shapes. Rather than from active bending (*7, 26*) or compressive buckling (*5, 6*), the deformations reported here arise from mechanically frustrated in-plane tension fields. These results establish a direct link between active nematic forces and morphogenesis, providing a mechanism for the rational design of programmable living materials.

## Results

### Emergent active forces in unconfined cellular nematics

To directly investigate how nematic forces drive tissue reshaping, we first sought to evaluate the forces and morphogenetic potential of fibroblast-based cellular nematics. To this end, we cultured monolayers of NIH-3T3 fibroblasts on soft elastomeric substrates (≈ 3 kPa) (*27*) coated with fluorescent tracers and the extracellular matrix protein fibronectin (Methods). As reported in previous studies (*28–30*), fibroblast monolayers display cellular alignment and self-organize long-range nematic order, which is lost locally at the cores of 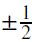 topological defects (Fig.1A and Methods). Labeling of F-actin, DNA, and phosphopaxilin revealed the presence of nematically aligned stress fibers, elongated nuclei, and widespread engagement of focal adhesions across the monolayer (Fig. S1 and Methods). Note that unlike turbulent-like cellular nematic systems (*10, 31*), topological defects in NIH-3T3 monolayers remained pinned at their formation sites (Movie S1). While individual cells were able to move along the nematic directions, the nematic field remained stable over time, presumably due to ECM deposition in a nematic state, thereby freezing the cellular nematic state (*30*). In contrast to jammed epithelial cell monolayers (*32*), which lack orientational order, fibroblast monolayers maintained a well-defined solid-like active nematic phase.

**Figure 1:**
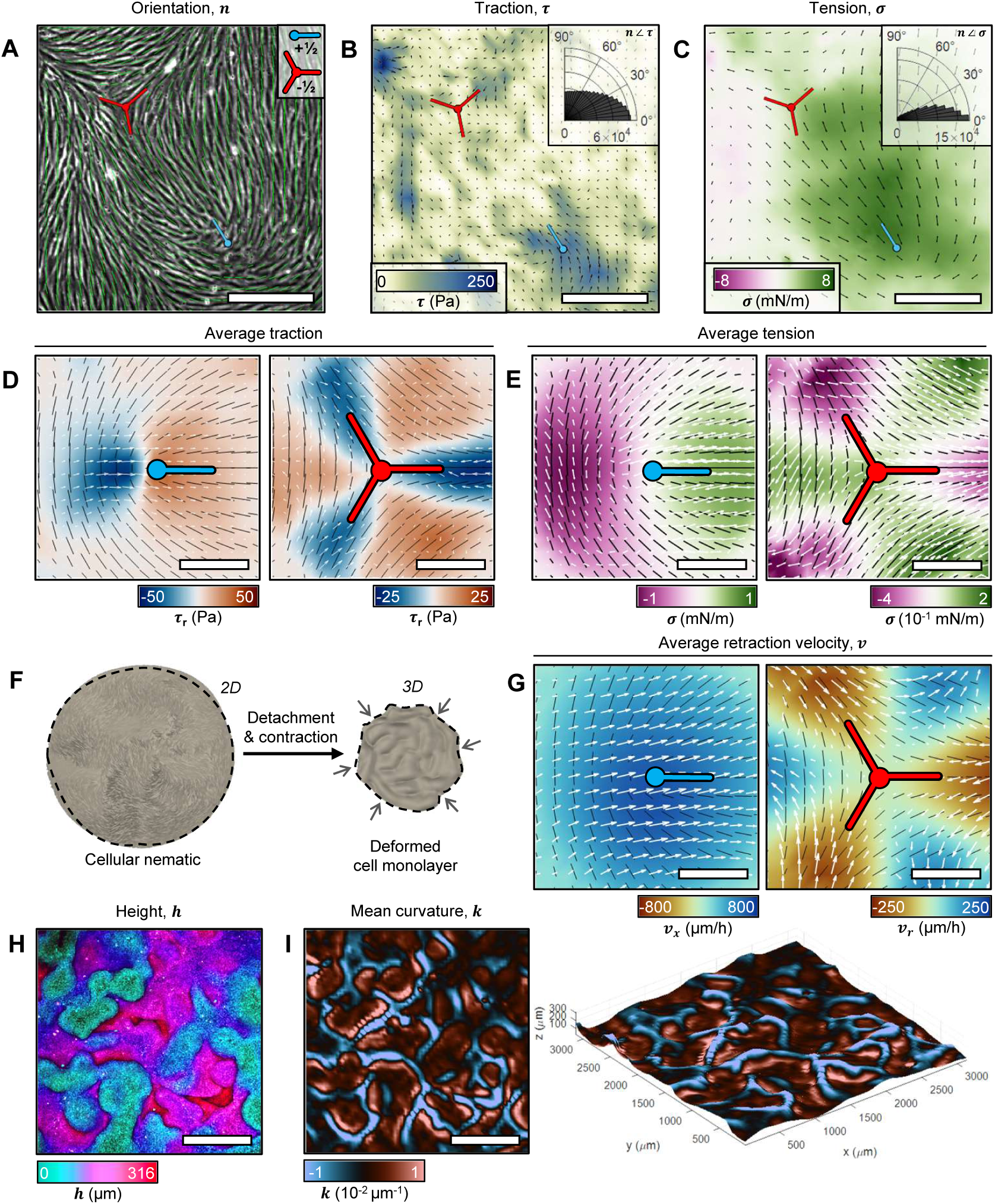
Morphology, active mechanics, detachment dynamics, and out-of-plane deformation of unconfined cellular nematics. (**A**) Phase contrast micrograph of a fibroblast monolayer with the corresponding orientation field (green vectors). The red and cyan symbols indicate the position of 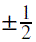 topological defects. (**B**) Traction field exerted by the monolayer in A. The color map represents the magnitude of the traction, the local directions of which are represented with black vectors. Green vectors indicate local orientation. The inset shows the distribution of angles between the orientation and traction vectors. (**C**) Tension field inferred from the traction field in B. The black vectors represent the local direction of the largest principal tension component, σ. The color map indicates the magnitude of the stress. Green vectors indicate local orientation. The inset shows the distribution of angles between the orientation vectors and the principal tension component. Data from N=15 positions with 30 time points each, spanning a duration of 10 h. (**D**) Average orientation and traction fields for 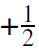 (left) and 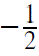 (right) topological defects. The black vectors represent the average orientation. The white vectors represent the tractions. The color map shows the magnitude of the radial component of the tractions. (**E**)Average orientation and tension fields for 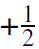 (left) and 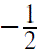 (right) topological defects. The black vectors represent the average orientation. The white vectors represent the local direction of the largest principal tension component. The color map shows the magnitude of the tension. The average fields were obtained with data that spanned a duration of 1 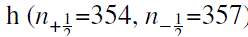. (**F**) Schematic illustrating the hypothesized reshaping of a cellular nematic during detachment. (**G**) Average velocity around 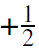 (left) and 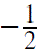 (right) topological defects right after monolayer detachment (N = 16 positions, 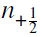 = 119, 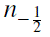 = 113). Color maps indicate the horizontal velocity component for 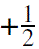 defects and the radial velocity component for 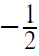 defects. (**H**) Z-projection of a membrane-stained free-floating monolayer of fibroblasts. The color map indicates the height *Z*. (**I**) Mean curvature (*k*) map corresponding to the confocal z-stack projected in panel H (top and 3D views). Scale bars for A-G: 100 μ*m*. Scale bars for H and I: 400 μ*m*.

Using traction force microscopy (*32*) (TFM, Methods), we first characterized the tractions exerted by cellular nematics on deformable substrates. High traction areas were observed in the vicinity of topological defect cores but vanished in aligned regions (Fig.1B). Globally, traction directions were correlated with cellular orientations (Fig. 1B, inset). The stress tensor within the tissue, inferred from the traction fields by using Monolayer Stress Microscopy (*33*) (MSM, Methods), revealed maximal tension between defects (Fig.1C), with its principal axis aligned with cellular orientation (Fig. 1C, inset). Averaging traction and tension fields over hundreds of defects revealed characteristic mechanical patterns (Fig.1D,E). In particular, 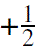 defects generated tractions following the head-to-tail direction, whereas 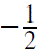 defects showed a three-fold symmetric pattern of balanced positive and negative radial tractions (Fig.1D). Despite substantial differences in magnitude, defect-mediated traction fields were qualitatively reproducible, particularly those generated by 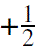 defects, as evidenced by the small variance of traction fields registered over ≈20h after confluency (Fig. S2A,B). Over time, the spatial distribution of defect-mediated traction fields did not change (Fig. S2C), but the traction magnitude increased (Fig. S2D), potentially due to cell proliferation and extracellular matrix remodeling (*34*). Similar to traction fields, defect-mediated tension fields were highly reproducible (Fig. 1E): 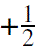 defects exhibited a highly tensile region at their tails and a compressive zone ahead of the defect core, whereas 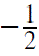 defects showed a three-fold symmetric tension pattern, with positive tension between the arms and compressive regions aligned along them, both fields consistent with observations in nematic epithelial monolayers (*10*). These results show that our cellular nematic sheets are in a state of internal tension patterns organized around nematic defects.

### Morphogenetic transformations in unconfined cellular nematics

We next hypothesized that, upon rapid detachment from the substrate, the cellular nematics would relax their active tension field by generating out-of-plane deformations (Fig.1F). To test this, we detached the tissues by enzymatically digesting the extracellular matrix with collagenase (Methods), thereby allowing them to contract freely. To prevent tissue fracture during this process (Movie S2), we inhibited tension with Blebbistatin, leading to the relaxation of the underlying elastic substrate (Movie S3, Fig. S3 and Methods). Despite this treatment, a residual basal level of contractility persisted (Fig. S3C). After the addition of Blebbistatin, the elimination of tissue-substrate adhesion triggered in-plane tissue contraction accompanied by rapid multicellular movements, aligned with nematically-guided tension fields (Fig. 1G, Movie S4). This transient dynamic state led to collective movements reminiscent of contractile active turbulence (*35, 36*). Shortly after this initial phase of in-plane dynamics, out-of-plane deformations emerged within minutes, resulting in the formation of wrinkled, free-floating cell monolayers (Fig. 1H,I). Laser ablation revealed that, rather than fully relaxing, these cell monolayers remained under residual tension, stored in elastically frustrated 3D configurations (Movie S5). These findings demonstrate that defect-mediated stresses in cellular nematics can drive rapid tissue rearrangements and morphogenetic transformations. We reasoned that, by controlling mechanical stress distributions, this process could be harnessed for effective shape programming.

### Control of cellular nematics through anisotropic micropatterns of adhesion

To control stresses in cellular nematics, we sought to guide cell alignment and impose specific nematic arrangements. This was achieved by chemically micropatterning substrates with anisotropic adhesive fibronectin patterns surrounded by a polyethylene glycol (PEG) cell-repellent coating, preserving a flat surface topography (Fig.2A and Methods). The micropatterns featured non-adhesive lines (≈ 2μ*m* in thickness) that guided the initial direction of cell spreading (*37*), setting preferred orientations prior to cell confluency (Fig.2B, Movie S6). Tissue growth progressed similarly to the non-patterned conditions, indicating that micropatterning did not alter growth dynamics (Fig. S4). After 2-3 days in culture, cellular nematics formed with topological defects in the positions predefined by the micropatterns (Fig. 2C-F). Although our approach can generate a wide range of cellular nematic configurations, we focused on minimal nematic arrangements with 2, 4, or 6 topological defects (Fig. 3 and Methods). While isotropic adhesive domains led to nematic arrangements with seemingly random spatial distributions (Fig.3A), anisotropic micropatterns promoted the formation of nematic monolayers with defects at specific locations, as shown by the average topological charge fields (Fig.3B) and the reproducible spatial distribution of cellular anisotropy and nematic order (Fig. S5, Methods). This micropatterning-based control strategy enabled robust and reproducible programming of cellular nematic orientation and defect positioning, while preserving compatibility with soft substrates for the probing of cellular forces.

**Figure 2:**
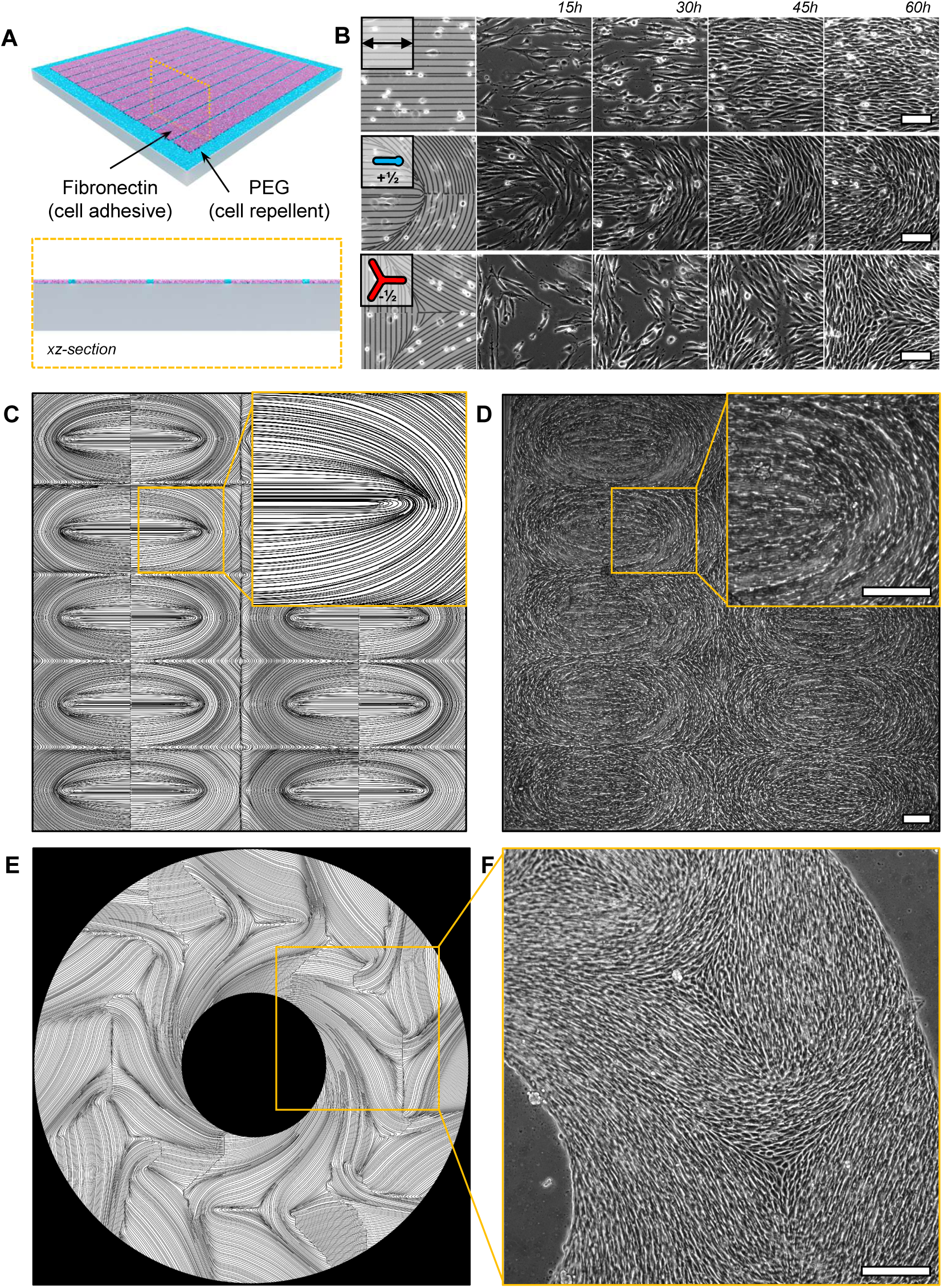
Control of cellular nematics using anisotropic adhesion micropatterns. (**A**) Schematic representation of an anisotropically micropatterned substrate. (**B**) Phase-contrast images of fibroblast monolayer regions evolving on micropatterns designed to impose different alignment constraints: uniform alignment (top), 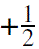 topological defect (middle) and 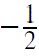 topological defect (bottom). The corresponding micropatterns are shown in the left panels. Scale bars: 100 μ*m*. (**C**) Anisotropic micropattern design featuring an array of 10 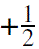 topological defect pairs. (**D**) Phase-contrast image of a fibroblast monolayer cultured on the micropattern shown in C for three days. (**E**) Anisotropic micropattern design featuring a disk with alternating 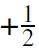 and 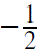 topological defects. (**F**) Phase-contrast image of a fibroblast monolayer cultured on the micropattern shown in E for three days. Scale bars: 200 μ*m*.

**Figure 3:**
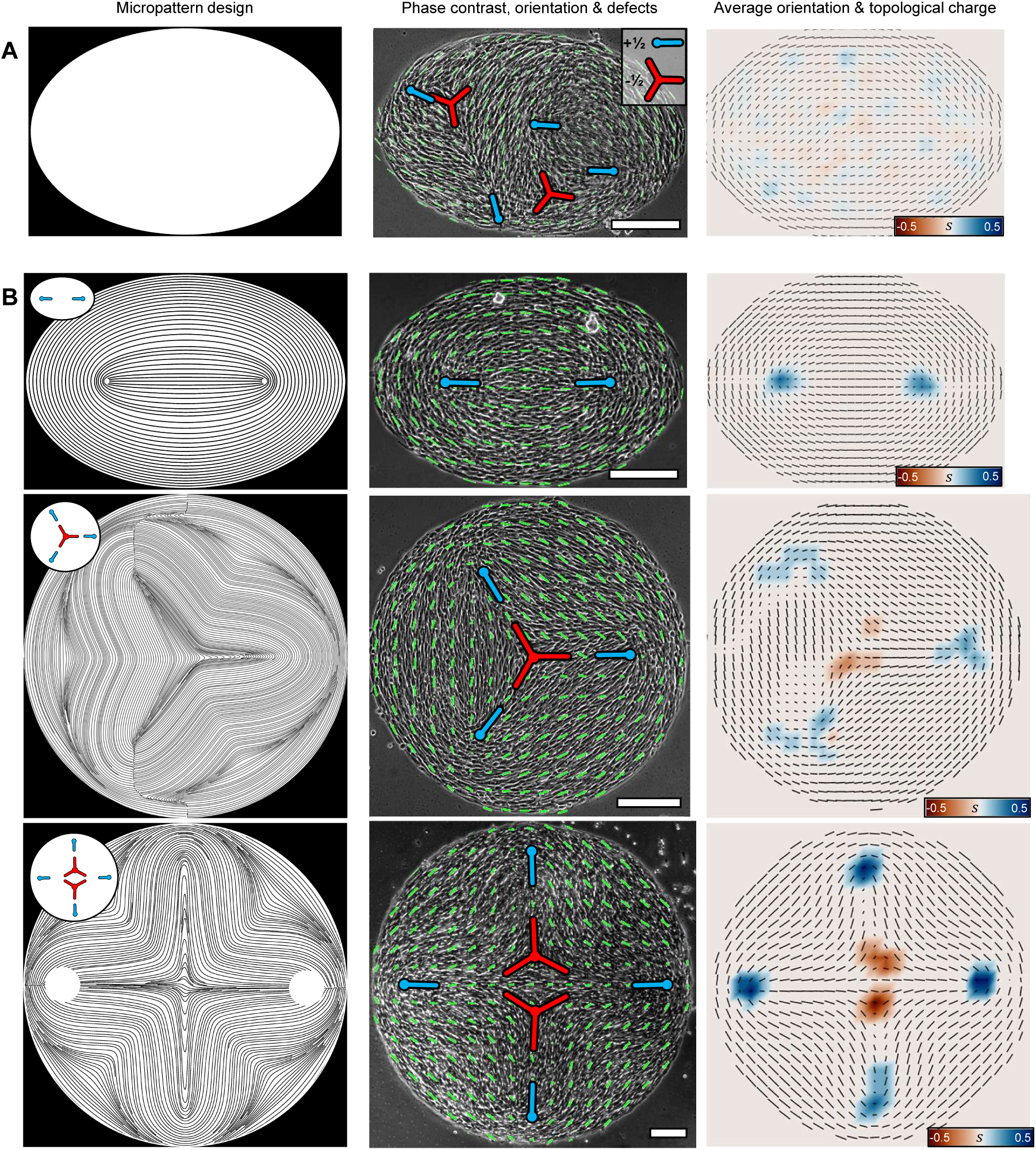
Minimal orientation-programmed cellular nematics. (**A**) Isotropic micropatterns of adhesion. From left to right: the micropattern designs (left column), phase contrast micrograph overlaid with local orientation vectors (green) and 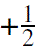 topological defect locations (central column), and average topological charge *s* fields (color map) with orientation fields (black vectors) (n = 22, right column). (**B**) Anisotropic adhesion micropatterns designed to impose, from top to bottom, 2-, 4-, and 6-topological defect arrangements (n = 29, 5, and 3, respectively). Columns as in A. Scale bars: 200 μ*m*.

### Pre-designed stress fields in cellular nematics by patterning topological defects

We next assessed the active mechanics of our controlled cellular nematics. To do this, we micropatterned the spatial arrangements comprising 2, 4, and 6 topological defects onto soft elastomeric substrates (Fig.4, Methods). Controlled cell alignment into precise orientational configurations led to very characteristic traction distributions dominated by the topological defects’ positioning (Fig.4C). The tractions around the topological defects were consistent with those observed in unconfined cellular nematics (Fig.1D): 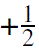 topological defects exerted pulling traction on the substrate along the head-to-tail direction and 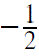 defects pulled inwards along the arms towards the core and pushed outward between arms. Next, from the obtained traction fields, we inferred monolayer tension (Methods). As in the unconfined case, the principal tension component aligned with cellular orientation (Fig.4D). For 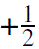 defects, we observed that the tension was minimal near the core and maximal behind the tail, consistent with their characteristic pulling traction behavior. Higher stresses at the center of the 2-defect monolayers were consistent with the accumulation of fibroblasts behind 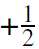 topological defects (Fig. S6), which can act as sources of multilayering (*38*). To confirm the tension build-up between 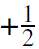 topological defects, we employed laser ablation to perform cuts between defect pairs, perpendicularly to cellular orientation, both at the center and boundary of the pattern. The monolayers retracted along the nematic direction, supporting that active tension is largely uniaxial along the nematic field as indicated by tension inference. Retraction speeds were similar regardless of the position of the cuts, although retractions near the boundaries were slightly shorter in range (Fig. S7A, Movie S7). Interestingly, however, only laser cuts made precisely between defect positions led to substrate relaxation under the topological defects (Fig. S7B, Movie S7), demonstrating that tension lines emerge between defects, mechanically linking them. Finally, we found that 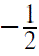 defects exhibited divergent tension fields. However, their characteristic three-fold symmetric compression-tension patterns were not observed, likely masked by the surrounding tension generated through interactions with neighboring 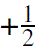 defects (Fig.4D). These experiments show the prominent effects of cellular orientation in the build-up of anisotropic stress in living tissues, with topological defects governing its spatial distribution.

**Figure 4:**
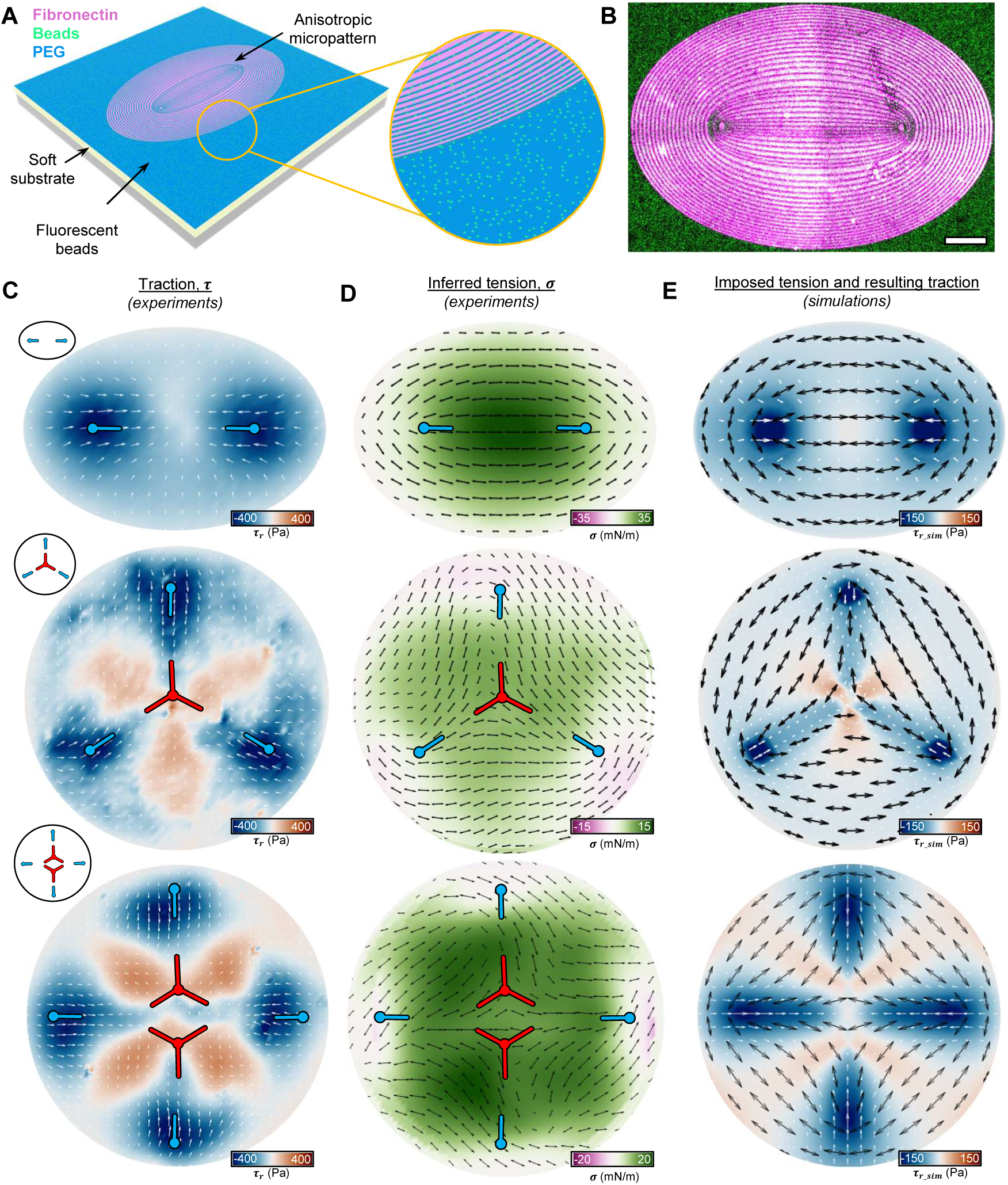
Programmed force fields in cellular nematics by harnessing topological defects. (**A**) Schematic representation of the experimental setup. (**B**) Epifluorescence micrograph of a fibronectin micropattern (magenta) on a soft substrate coated with fluorescent beads (green). Scale bar: 100 μ*m*. (**C**) Average spatial distributions of traction. Colormap indicates the radial component of traction. (**D**) Average spatial distributions of tension for the 2-, 4-, and 6-topological defect arrangements, shown from top to bottom (n = 29, 5, and 3, respectively). The tension fields are represented by double black arrows and a colormap, indicating the directions and magnitude of the principal tension component, respectively. (**E**) Imposed tension fields in simulations (black arrows) and resulting traction fields for 2-, 4-, and 6-topological defect arrangements. Colormap indicates the radial component of tractions.

To understand the origin of the observed mechanics, we sought to model these minimal cellular nematic configurations *in silico*. Briefly, we first generated computationally 2D nematic fields with pre-defined defect positions akin to those observed experimentally. Consistent with the observations that cellular nematic patterns became frozen and developed anisotropic contractile tension along the nematic field, we modeled the system as a 3D nonlinearly elastic substrate fixed at the bottom and subjected to a 2D uniaxial and contractile tension along a nematic field imprinted on the free surface of the material (Fig, 4E, Supplementary Text). We did not attempt to fine-tune the thickness-dependent magnitude of active tension (Fig. S6) and instead considered it to be homogeneous. These simulations reproduced the experimentally observed traction patterns both qualitatively and quantitatively, confirming that nematic tension fields govern the mechanical behavior of these systems. Together, our experiments and simulations show how imposed inter-defect interactions lead to very characteristic force fields that emerge from mechanical defect-defect interactions, demonstrating the potential of micropatterning cellular orientation in finely controlling force distributions in nematic tissues.

### 3D reshaping of minimal cellular nematics

We next sought to leverage our simulations to predict how the interplay between nematic order and contractile forces gives rise to curvature. To this end, we assumed that, when detached, the nematic cell sheet remains in a solid state with frozen nematic texture attached to the material, and modeled it as a thin elastic shell subjected to a uniaxial and contractile tension along the nematic field (Supplementary Text). This model reproduced previous results about the reshaping of thin sheets made of responsive and patterned liquid crystal elastomers (*39–41*) or baromorphs (*42*). We simplified the detachment process by simulating the mechanical relaxation of the pre-stressed shell embedded in a viscous environment, where the active tension was ramped and then held constant until the system reached a steady-state.

We began by modeling the deformation of elliptical sheets triggered by the contraction of minimal contractile nematics comprising two 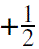 defects. As tension increased, thin sheets exhibited in-plane contraction along the axis connecting the 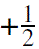 topological defects, following the direction of contractile stress (Fig.5A). As a consequence of contraction, the sheets experienced localized compression, triggering out-of-plane buckling and subsequent edge folding, ultimately resulting in a bowl-shaped morphology (Fig.5A and Movie S8). During this reshaping process, the initially flat monolayer adopted a doubly curved geometry. This is an instance of Gaussian morphing, in which a prescribed pattern of in-plane tensions cannot fully relax through in-plane deformation alone and, instead, the system buckles into the third dimension to minimize its overall elastic energy (*39, 43*).

**Figure 5:**
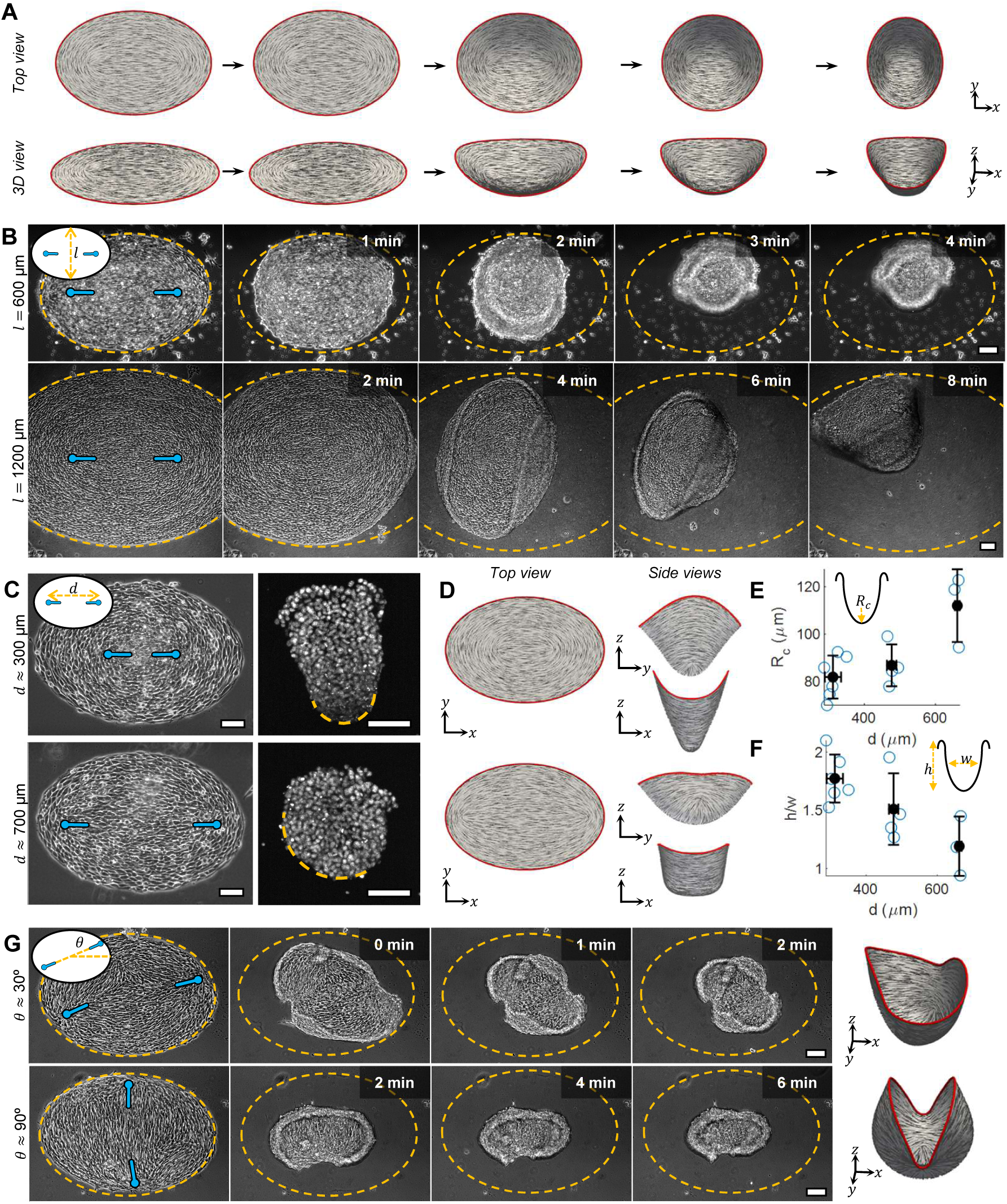
Control of tissue shape by harnessing two-defect cellular nematics. (**A**) Snapshots from simulations showing the deformation of 2-defect cellular nematics (top and orthographic views). Vectors depict orientation. Colormap indicates the order parameter. (**B**) Phase-contrast micrographs of detaching 2-defect cellular nematics with different sizes, both leading to a bowl-shaped configuration after detachment. The monolayer was treated with Blebbistatin before detachment. The time elapsed since the start of retraction is indicated. The dashed line indicates the perimeter of the monolayer at t=0. (**C**) Phase contrast micrographs of 2-defect cellular nematics with different inter-defect distances, and corresponding maximum Z-projections DNA-stained 3D tissue structures, obtained after their detachment. Dashed line indicates the base of the bowl-shaped morphologies. (**D**) Snapshots from simulations of post-detachment steady-state configuration of thin elastic shells exhibiting 2-defects nematic arrangements with different inter-defect distance (bottom and side views). (**E**) Minimum radius of curvature (*R*_*C*_) of 3D structures obtained from Blebbistatin-treated 2-defect cellular nematics with respect to the distance between 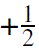 topological defects (*d*). (**F**) Height-width ratio (ℎ/*w*) of the same 3D structures with respect to the inter-defect distance (*d*). (**G**) Phase contrast time lapses of reshaping cellular nematics featuring pairs of 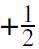 topological defects at 45° and 90° with respect to the long axis of the elliptical confining geometry. The right panels show the post-detachment steady-state configuration of thin elastic shells exhibiting the same initial nematic arrangements as in the experiments (orthographic views). Scale bars: 100 μm.

We then tested these predictions by enzymatically detaching similar cellular nematics. Consistent with our simulations, the detachment of 2-defect cellular nematics led to a rapid in-plane contraction along the axis connecting the defects, guided by the directions of maximal tensile stress (Fig.5B). This phase of defect-driven retraction was consistent with the characteristic early dynamics of detached 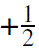 topological defects in experiments (Fig. 1G). Contraction was accompanied by a gradual irreversible loss of cellular anisotropy (Fig. S8). Subsequently, as captured by our simulations, the retraction of the monolayers led to buckling in the radial direction and to folding into a bowl-shaped geometry (Fig.5B, Movie S9 and Methods). In simulations, spatial variations in the sheets’ thickness had a negligible effect on the resulting shapes (Movie S10), indicating that shape selection is primarily governed by the orientations of nematic tension fields.

### 3D tissue reshaping through rational design of nematic order

We next explored the potential of geometrical parameters and defect positioning to control 3D tissue reshaping, combining both experiments and simulations. Systems featuring the same nematic arrangement but twice the area, i.e. increased slenderness, exhibited similar reshaping dynamics (Fig. 5B and Movies S11 and S12). The inter-defect distance prior to detachment significantly influenced the resulting tension fields (fig. S9), and consequently, the morphology of the emergent 3D structures (Fig. 5C–F; Movies S13–S16). The angle between the inter-defect axis and the long axis of the elliptical confinement also influenced the reshaping dynamics and final morphology (Fig. 5G and Movies S17-S20). Finally, the detachment of randomly arranged defect configurations with identical elliptical geometries resulted in markedly different 3D tissue morphologies, discarding major effects from the geometry of the cellular nematics (Movie S21). In all these cases, in-plane contraction and subsequent folding were initiated by topological defects’ retraction. The reshaping dynamics and resulting morphologies were governed by anisotropic tension fields, intrinsically linked to the nematic order and the spatial arrangement of topological defects.

Our experiments and simulations demonstrate that the guidance of nematic order enables the formation of self-shaping active layers with a well-defined tension distribution, governed by the interplay between nematic order and contractility. To further explore the potential of this framework, we investigated the generation of more complex structures by reshaping circular cellular nematics comprising 4 or 6 topological defects. As in the 2-defect configuration, we observed an initial retraction of the tissue, driven by the nematic tension field, predominantly along the head-to-tail direction of the 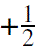 topological defects. Both in experiments and simulations this led to transient triangular monolayers in the case of the 4-defect configuration (Fig.6A and Movies S22 and S23) and squared monolayers in the case of the 6-defect configuration (Fig.6B and Movies S24 and S25). In both cases, the initial in-plane compression between defects triggered upward folding, producing complex bowl-shaped morphologies with regions of high curvature emerging at the initial sites of defect retraction. The resulting 3D structures exhibited a three-fold symmetry for the 4-defect case and a four-fold symmetry for the 6-defect case.

**Figure 6:**
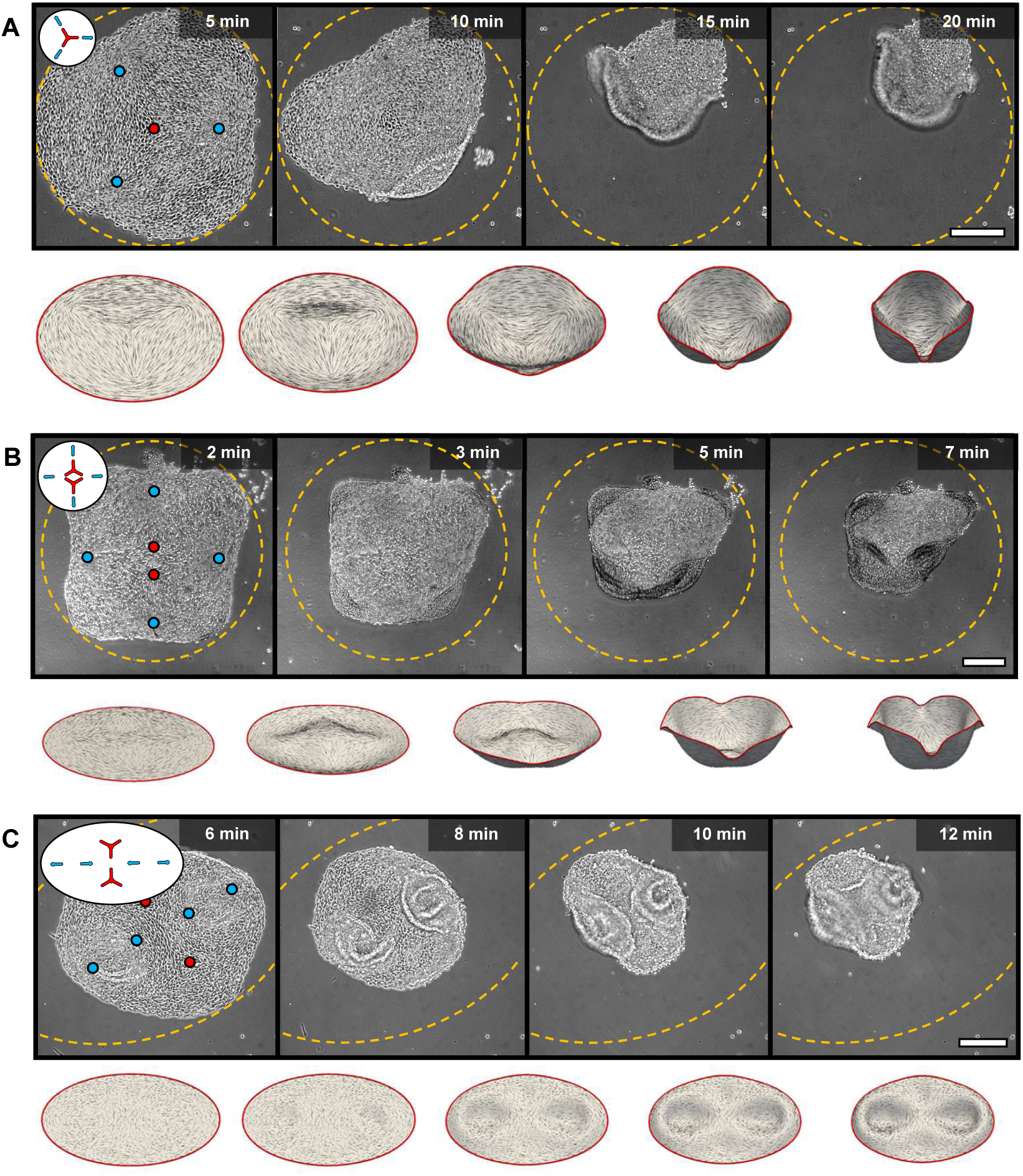
Hierarchical complexity in tissue shape driven by multiple interdefect interactions. Phase-contrast time-lapse images and simulation snapshots showing the reshaping of cellular nematics with 4-defect (**A**) and 6-defect (**B,C**) arrangements after detachment. The time elapsed since the initiation of collagenase treatment is indicated. The dashed line indicates the perimeter of the monolayers at t=0. Scale bars: 250 μ*m*.

Finally, we investigated the reshaping dynamics of an alternative 6-defect configuration, displaying head-to-head oriented 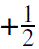 topological defects, which led to localized compressive stress (Fig.6C, fig. S10). Initial in-plane retraction was primarily driven by the tense regions between pairs of tail-to-tail oriented 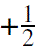 defects, which led to the formation of two distinct bowl-shaped invaginations (Fig.6C, Movie S26). In contrast, the compressive zone located between the head-to-head 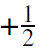 defects expanded in the direction connecting 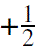 defects and contracted perpendicular to it, leading to a tissue morphology combining bowl-shaped invaginations and a saddle-shaped region (Fig.7, and Movie S27). In simulations, this configuration was metastable (Movie S28), indicating that additional factors present in the experimental system, such as extracellular matrix components, may contribute to stabilizing tissue shapes that would otherwise evolve further in a purely elastic model. Altogether, these experiments and simulations establish the coupling between nematic order and contractility as a fundamental mechanism for guiding tissue morphogenesis, also setting the ground for the rational design of tissue shape.

**Figure 7:**
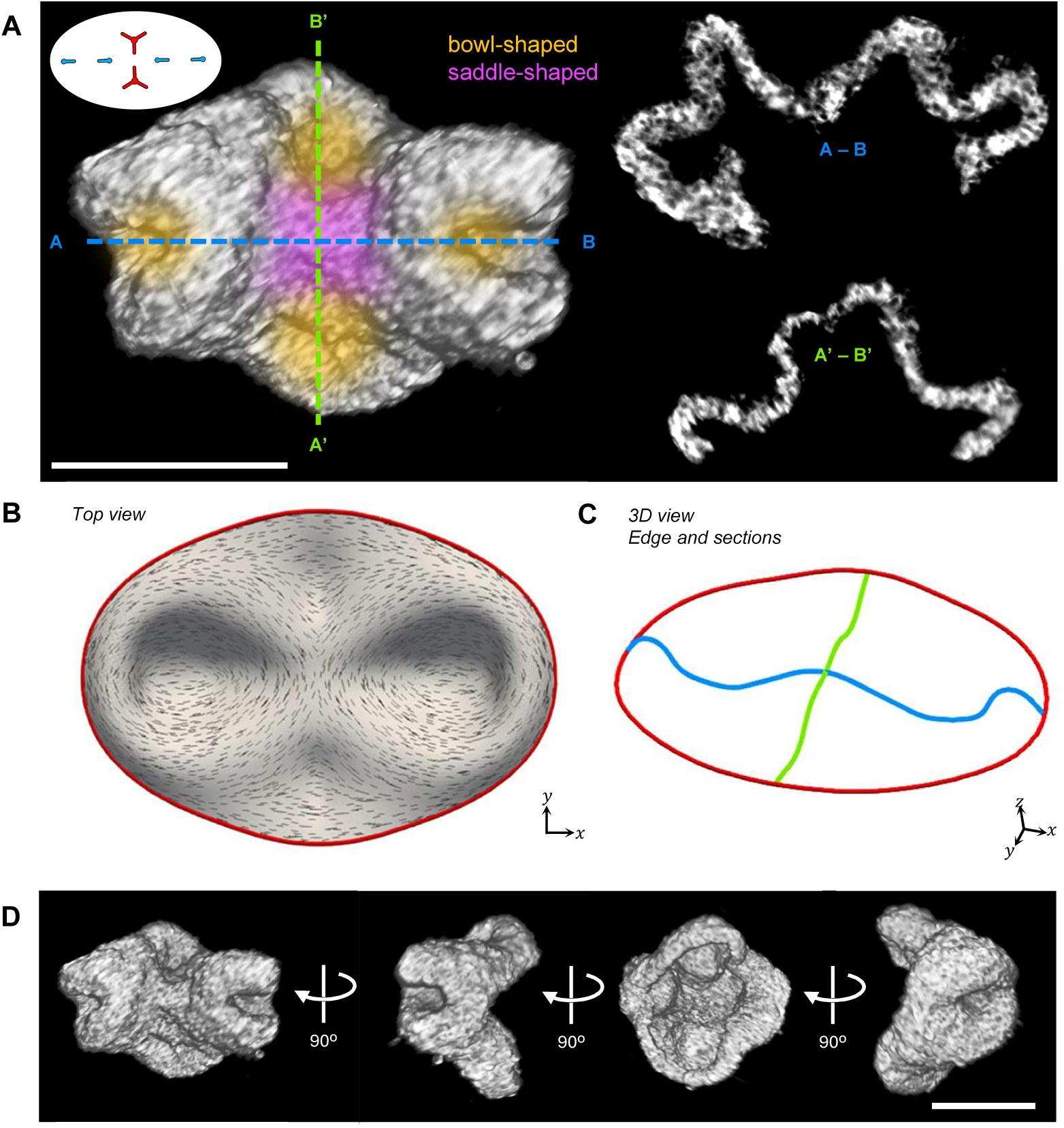
Three-dimensional tissue architecture shaped by high-order inter-defect interactions. (**A**) 3D rendering and orthogonal cross-sections of a tissue structure resulting from a 6-defect nematic configuration. 3D image stacks were acquired using light-sheet microscopy. (**B**) Snapshot from simulations of a deformed thin elastic shell exhibiting the same nematic arrangement as in A. (**C**) Tissue edge contour (red line) and tissue profile sections along the short and long axes (blue and green lines, respectively) corresponding to the shape in B. (**D**) Four 3D renderings of the same structure shown in A, each rotated by 90°. Scale bars: 250 μ*m*.

### Conclusions and outlook

Our findings reveal novel emergent behaviors in cellular tissues exhibiting nematic order, where the nematically guided tensions drive dynamic shape transformations toward specific morphologies. We show that the morphing principle operating in our system relies on mechanically frustrated in-plane tension patterns imprinted on a thin solid surface. This contrasts with previous conceptual frameworks for reshaping of active nematic surfaces, considering dynamical nematic fields evolving on fluid surfaces (*18–23*), and relying on physical mechanisms not present in our model such as active flows, Frank free-energy relaxation (*19, 44*), or active torques (*20*).

More generally, reshaping of animal cellular sheets is generally attributed to active bending (*7, 26*) or compressive buckling (*5, 6*). Instead, the principle reported here is closely related to a prevalent morphing principle in plant tissues such as leaves and flowers, namely Gaussian morphing, according to which, out-of-plane reshaping is induced by geometric frustration of incompatible in-plane growth patterns (*43, 45–48*). Here, the geometric incompatibility is generated by nematic tension patterns rather than growth (Supplementary Text). Due to the anisotropy of tension along a texture of orientations, our system is a living and microscopic analog of artificial implementations of Gaussian morphing in striated surfaces such as liquid crystal elastomer sheets (*39–41*) or baromorphs (*42, 49*).

Orientational order and topological defects are increasingly identified in living tissues, both during development (*16, 17, 50, 51*) and disease (*52–54*), highlighting the natural relevance of nematic organization in biological systems. In this context, our findings suggest that nematically guided Gaussian morphing may represent an active mechanism contributing to tissue remodeling processes *in vivo*. Taken together, the strategies presented here provide a robust toolkit for investigating defect-mediated mechanical interactions and shape emergence in nematic tissues, while establishing the foundation for harnessing cellular nematics as mechanically encoded, shape-programmable living materials.

## Acknowledgments

We thank the members of the Trepat, Arroyo, and Roca-Cusachs groups for their insightful comments on this work. We are grateful to O°. O° zgüç (Institute for Bioengineering of Catalonia, IBEC, Barcelona) for guidance on the fixation of 3D tissue structures, and to L. Bardia and J. Colombelli (Advanced Digital Microscopy core facility, Institute for Research in Biomedicine, IRB, Barcelona) for their assistance with the mounting, clearing and imaging of 3D tissues.

## Funding

P.G. acknowledges support from the Horizon Europe Research and Innovation Program of the European Union through Marie Sklodowska-Curie Actions (grant agreement 101065794). P.R.-C. acknowledges funding from the Spanish Ministry of Science and Innovation (PID2022-142672NB-I00), the European Research Council (AdG 101097753), the Generalitat de Catalunya (2017-SGR-1602), and the prize ‘ICREA Academia’ for excellence in research. M.A. acknowledges funding from the Spanish Ministry of Science and Innovation (PID2022-142178NB-I00), and the Generalitat de Catalunya for the prize ‘ICREA Academia’ for excellence in research. X.T. acknowledges funding from the Generalitat de Catalunya (AGAUR SGR-2017-01602), the CERCA Programme, the Spanish Ministry for Science and Innovation MICCINN/FEDER (PID2021-128635NB-I00 MCIN/AEI/10.13039/501100011033 and “ERDF-EU A way of making Europe”), European Research Council (AdG 883739), Fundació la Marató de TV3 (project 201903-30-31-32), European Commission (H2020-FETPROACT-01-2016-731957), La Caixa Foundation (LCF/PR/HR24/00326) and the Human Frontiers Science Program (HFSPRGP022/2024); IBEC is recipient of a Severo Ochoa Award of Excellence from the MINECO.

## Author contributions

P.G. and X.T. conceptualized the research. P.G. designed and performed the experiments and analyzed the data with the collaboration from M.G.-G.; W.M. and P.K.B performed the numerical simulations and wrote the Supplementary Text; X.T. and M.A. supervised the research; P.G. wrote the manuscript with the collaboration from all the authors.

## Competing interests

The authors declare no competing interests.

## Data and materials availability

The raw data will be made available by the corresponding authors through an online repository.

## Supplementary materials

Materials and Methods Supplementary Text Figs. S1 to S10 Movies S1 to S28

## Notes

### Competing Interest Statement

The authors have declared no competing interest.

