## Supplementary Material for "Guidance of cellular nematics into shape-programmable living surfaces"

### Supplementary Materials for
### Guidance of cellular nematics into shape-programmable living surfaces

Pau Guillamat[1,2*], Waleed Mirza[3†], Pradeep K. Bal[4†], Manuel Gómez-González[1,2],
Pere Roca-Cusachs[1,2], Marino Arroyo[4*] & Xavier Trepat[1,2*]
[1]Institute for Bioengineering of Catalonia (IBEC), Barcelona, 08028, Spain.
[2]The Barcelona Institute for Science and Technology (BIST), Barcelona, 08028, Spain.
[3]European Molecular Biology Laboratory (EMBL), Barcelona, 08003, Spain.
[4]LaCàN, Universitat Politècnica de Catalunya (UPC) – BarcelonaTech, Barcelona, 08034, Spain.
*Corresponding authors.

†These authors contributed equally to this work.

**This PDF file includes:**

Materials and Methods

Supplementary Text

Figs. S1 to S10

Captions for Movies S1 to S28

**Other Supplementary Materials for this manuscript:**

Movies S1 to S28

### Materials and Methods

#### Experimental techniques

##### Cell culture

NIH/3T3 mouse fibroblasts were cultured in DMEM media containing 4500 mg/L glucose, 1mM sodium pyruvate (Life Technologies) and supplemented with 10% fetal bovine serum (FBS), 100 units/mL penicillin and $100\mu g/mL$ streptomycin. Cells were maintained at 37ºC under 5% $CO_2$ and were not allowed to become confluent. The maximum passages were kept below 20.

##### Preparation of soft substrates

To prepare soft elastomeric surfaces (≈3 kPa) (*27*), glass bottom dishes (35-mm, no. 0 coverslip thickness, Mattek) were first spin-coated (90 s at 400 rpm) with a $\approx 100\mu m$ layer of a 1:1 mixture (weight ratio) of CY52-276A and CY52-276B polydimethylsiloxane (Dow Corning Toray), previously degassed for 30 minutes on ice. The dishes were then cured at 70°C overnight in an oven. The substrates were kept at room temperature in a clean and dry environment and used within 2 months after fabrication.

##### Coating of soft substrates with fluorescent tracers

After curing, a polydimethylsiloxane (PDMS) stencil (≈250 $\mu m$ thick) with an inner diameter of 7mm was placed on top of the soft substrates. The inner region of the stencil was treated with (3-aminopropyl)triethoxysilane (APTES, Sigma-Aldrich, cat. no. A3648) diluted at 5%v/v in absolute ethanol for 3 min, rinsed 3 times with absolute ethanol and rinsed once with ultra pure water. The samples were incubated for 1 h with a filtered (450 nm filter) and 10-min sonicated solution of red fluorescent carboxylate modified beads (FluoSpheres, Invitrogen) of 200 nm in diameter in borate buffer (3.8 mg/ml sodium tetraborate, Sigma-Aldrich and 5 mg/ml boric acid, Sigma-Aldrich). The gels were then rinsed 3 times with borate buffer.

**Passivation of soft substrates**

Prior micropatterining, the soft surfaces were treated with a 0.1 mg / ml polylysine solution (PLL, Sigma P2636) in ultra-pure water for 1 h, then washed with 10mM HEPES buffer (pH=8.4). A solution of 50mg/mL mPEG (MW 5,000) - Succinimidyl Valerate (SVA) (Laysan Bio) in the same HEPES buffer was applied to passivate the surface for 1.5 h and then washed out with phosphate buffer saline (PBS). The substrates were stored at 4 ºC and normally used within 48 h after passivation.

**Micropatterning of soft substrates**

Micropatterns were generated by using a UV-activated mPEG-scission reaction, spatially controlled with the PRIMO system (Alvéole) (*55*), mounted on a Nikon Eclipse Ti-e inverted microscope. In the presence of a photoinitiator compound (PLPP, Alvéole), the antifouling properties of the PEGylated substrate are tuned by exposure to patterns of near-UV light ($\lambda$=375nm). Patterns were designed either with Matlab (MathWorks, Inc.) or with ParaView, considering the morphological features of our unconfined cellular nematics. After illumination (900 mJ/mm$^2$) through a Plan Fluor x10 objective (NA = 0.3), PLL regions were exposed. After rinsing with PBS, fibronectin (SIGMA, F0895) was incubated at 100 $\mu g/mL$ at room temperature for 5 minutes to coat the PEG-free regions with the cell-adhesive protein. In some cases, AlexaFluor-647-conjugated fibrinogen or fibronectin was added to the previous fibronectin solution to allow visualization of micropatterns. The excess protein was washed out with PBS. The patterned substrates were stored at 4ºC and used within 48 h after preparation.

**Cell seeding**

Before cell seeding, PBS was replaced by medium and a cell suspension was added at densities of $\approx 10^5$ cells/cm$^2$. The samples were kept in an incubator at 37 ºC and 5% $CO_2$. After 15 min, non-adhered cells were washed and 2 ml of medium were added. Cells were seeded between 2-3 days prior to the experiments.

##### Time-lapse imaging

Time-lapse imaging was performed with an inverted Nikon Ti-E microscope installed in a thermostatically controlled chamber (Life Imaging Services) and equipped with a microincubator for $CO_2$ and humidity control (The Brick, Life Imaging Services). The microscope was also equipped with an automated stage and a Yokogawa CSU-W1 spinning disk unit. Image acquisition was performed with an Andor Zyla 4.2 Plus camera, operated with Micro-Manager Software. We performed phase contrast (10/20x objectives, NA 0.3/0.45). Typically, we acquired 12 images/h for at least 10h.

##### Laser ablation and imaging

Laser ablation was performed through a Plan Fluor x10 objective (NA = 0.3) using the Rapp OptoElectronic UGA-42 / Z Firefly point scanning device, coupled to an inverted Nikon Ti-2 microscope. During ablation, samples were mantained at 37 ºC and in a micro-incubator for $CO_2$ and humidity control. During ablation, we imaged our samples in both bright-field and epifluorescence modes, acquiring time-lapses at a rate of 4 images/s. The retraction dynamics was characterized by Particle Image Velocimetry (PIV), using the PIVlab App (*56*) in Matlab (MathWorks, Inc.).

##### Inhibition of contractility

For inhibiting contractility (myosin-II ATPase), cells were treated with para-NitroBlebbistatin (Motorpharma) at 10 $\mu M$, keeping the DMSO concentration below $10^{-3}$%v/v.

##### Detachment of cell monolayers

To detach cell monolayers from their substrates we used collagenase (SIGMA, C2674-100MG) at a final concentration of 500 U/mL. Collagenase aliquots were prepared in Hanks balanced salt solution (HBSS) at a concentration of 20 kU/mL and were stored at -20 ºC.

##### Fixation of monolayers, fluorescence labeling and imaging

Cells were fixed with 4% paraformaldehyde (Sigma) at 37ºC dissolved in PBS for 10 minutes.

Actin was labeled with SPY-actin-555 (Spirochrome). Concentrations used were $1\mu M$ (30 minutes incubation). Cell nuclei were labeled after fixation with Hoechst 33342 (Thermofischer) at $10\mu g/mL$ (5 minutes incubation). Membranes were stained with Wheat Germ Agglutinin (WGA, Invitrogen) conjugated with Alexa Fluor 488 at $10\mu g/mL$ (5 minutes incubation). For immunostaining of Phospho-Paxilin, we used the primary antibody Phospho-Paxillin (Tyr118) #2541 (Cell Signaling Technology).

Fixed samples were imaged using a Nikon Eclipse Ti-E microscope equipped with a Yokogawa CSU-W1 spinning disk confocal unit. We used air objectives x10 (NA 0.3), x20 (NA 0.75) and x40 (NA 0.75). The microscope was operated with Micro-Manager software.

**Fixation, staining, mounting, clearing and imaging of 3D tissue structures**

3D tissue structures were aspirated with a micropipette and introduced into droplets containing 4% paraformaldehyde (Sigma), placed under a layer of silicone oil (Bluesil, viscosity 0.02 Pa·s, BlueStar Silicones) in cell culture dishes, 35mm in diameter. After 10 minutes, tissues were retrieved from the paraformaldehyde droplets and suspended in PBS. To rinse the tissues, half the PBS volume was exchanged five times.

For cell membrane staining, half the PBS volume was replaced with a 1:100 dilution of a Wheat Germ Agglutinin solution (ThermoFisher) in PBS. Tissues were incubated for 5 minutes at room temperature. After staining, tissues were rinsed by exchanging half their surrounding volume with PBS five additional times. Fixed and stained 3D tissue structures were stored in PBS at 4°C until mounting.

Before imaging, fixed 3D tissue structures were suspended in a low-melting-point agarose solution ($0.7\% w/v$). A 1 mL syringe was used as a mold for the casting. After the agarose solidified, cylindrical casts were carefully removed from the syringe mold and stored in PBS at 4°C until further use.

For improved imaging, tissues embedded in agarose blocks were cleared by immersion in OptiPrep solution (Sigma, D1556) under gentle agitation on a rocker for 24 hours.

3D tissue structures were imaged using a custom-built Selective Plane Illumination Microscope (MacroSPIM), based on Nikon AZ100M macroscopes with dual-view detection and dual-side illumination. Light-sheet formation was achieved using a cylindrical achromatic lens with a 50 mm focal length, producing a light-sheet waist of approximately 3–4 $\mu m^2$ at 488 nm (diode laser excitation). Samples were mounted in a quartz cuvette filled with OptiPrep solution (Sigma, D1556). Imaging was performed using a 525/50 nm bandpass emission filter at an effective zoom of 9.6× (2× objective lens with internal 8× zoom and a 0.6× tube lens), yielding a lateral pixel size of 677 nm. Axial sampling was conducted with a Z-step of 2.5 $\mu m^2$. To minimize striping artifacts, light-sheet pivoting at 150 Hz was implemented via a galvanometric scanner. For the dataset shown in Fig. 7, single-view and single-side illumination were used. 3D image stacks were rendered using Imaris Viewer (OXFORD Instruments). Selected cross-sections were generated using the Ortho Slicer tool.

#### Image processing and analysis

##### Analysis of cellular orientation

The local orientation field $\hat{n}(x, y)$ in cell monolayers was extracted by using the plugin OrientationJ in Fiji (*57*), which is based on the structure tensor method (*58*). In summary, for each pixel of an image, intensity gradients are computed. A structure matrix is then obtained from the products of the components of the intensity gradients that are averaged in local sub-windows. For most of the cases, the sub-window size was set to 6x6 $\mu m^2$. Diagonalization of this structure matrix results in two eigenvectors per pixel. The eigenvector with the smallest eigenvalue represented the direction of the smallest variation of the intensity map in the vicinity of the pixel and was associated with the main orientation $\theta$, the local orientation of $\hat{n}$ with respect to a fixed axis. The amplitude of the orientation vectors, referred to as Anisotropy in the text, was also extracted from OrientationJ, where it is referred to as Coherency, $C$. Finally, the values of $\theta$ and $C$ were then typically averaged in image sub-windows of 12x12 $\mu m^2$. For further details on the mathematical framework of this method, see (*58*). Further analyses from orientation data were performed with custom-written codes in Matlab (MathWorks, Inc.).

**Detection, classification and orientation of topological defects**

For the detection of topological defects, we build on previously used algorithms. First, we define as defect areas the regions where the order parameter $Q = \sqrt{\langle\cos 2\theta\rangle^2 + \langle\sin 2\theta\rangle^2}$ was below a threshold value of 0.3. The brackets $\langle\rangle$ denote spatial averaging in local regions. To assess the topological charge of defects, with topological charges $s = \pm\frac{1}{2}$, we calculated the winding number $\frac{\sum\Delta\theta}{2\pi}$, where $\sum\Delta\theta$ is the accumulated rotation of the orientational field around these low-order regions (*59*). The orientation of the topological defects with respect to a fixed axis, $\psi$, was calculated using the method in reference (*60*) as

$$\psi_{+\frac{1}{2}} = +\arctan\left[\frac{\delta_x Q_{xy} - \delta_y Q_{xx}}{\delta_x Q_{xx} + \delta_y Q_{xy}}\right] \tag{S1}$$

$$\psi_{-\frac{1}{2}} = -\frac{1}{3}\arctan\left[\frac{-\delta_x Q_{xy} - \delta_y Q_{xx}}{\delta_x Q_{xx} - \delta_y Q_{xy}}\right] \tag{S2}$$

where, $Q_{xx} = 2C\cos(2\theta)$ and $Q_{xy} = 2C\sin(2\theta)$. $C$ corresponds to the Coherency and $\theta$ to the local orientation of $\hat{n}$ with respect to a fixed axis. Both quantities, $C$ and $\theta$, were extracted with the OrientationJ plugin. The gradients of $Q_{xx}$ and $Q_{xy}$ were calculated using the function $gradient$ in Matlab (MathWorks, Inc.).

**Characterization of the cell monolayers' morphology and dynamics**

Surface curvature (Fig.1H-I) was calculated and mapped in Matlab (MathWorks, Inc.) using custom-made codes, including the function $surfature$.

To characterize the height profiles for 2-defect configurations (Fig. S6), we first obtained YZ-orthogonal cuts for ≈100 x positions between topological defects using the function $Reslice$ in Fiji. Second, we generated average intensity x-projections of the YZ-orthogonal cuts for each monolayer (N=8). Third, we manually extracted the coordinates of each x-projection using the $Multi-point$ selection tool in Fiji.

To characterize the dynamics of the monolayers, we performed tracer-free velocimetry analysis with a public domain Particle Image Velocimetry (PIV) plugin (*61*) in Fiji. In particular, we used iterative PIV with 3 passes, with a decreasing interrogation window size. The minimum window sizes used in the analysis were generally set at 10x10 $\mu m^2$, which showed good qualitative agreement with our experimental observations.

**Traction Force Microscopy**

Fluorescent beads were imaged before and after the detachment of the monolayers with trypsin or collagenase treatments. The 2D displacement of the upper surface of the PDMS substrate was then computed by comparing the reference image (without cells) with the image of each deformed configuration (with cells), through a custom Particle Image Velocimetry (PIV) implementation in Matlab (MathWorks, Inc.) (*62*). The 2D traction field, $\vec{\tau}(x, y)$ exerted by fibroblasts on the upper surface of our substrate, was then calculated by Finite Thickness Fourier transform Traction Force Microscopy (TFM) (*32, 63*).

**Monolayer Stress Microscopy**

The spatial distribution of in-plane internal stresses, $\vec{\sigma}(x, y)$, within the fibroblast monolayers was calculated from the cell-exerted traction field by solving the two-dimensional force balance equations using standard Monolayer Stress Microscopy (MSM) algorithms (*33*). MSM was implemented in Python 3 using the following libraries: NumPy (*64*), SciPy (*65*), Matplotlib (*66*), scikit-image (*67*), pandas (*68*), pyFFTW (*69*), opencv (*70*) and cython (*71*).

**Analysis of average fields around topological defects**

Average vector fields, including $\hat{n}$, $\vec{v}$, $\vec{\tau}$ and $\vec{\sigma}$, around $\pm\frac{1}{2}$ topological defects, were obtained by cropping vector field regions of a given size centered around the cores of the topological defects, which were rotated by $\psi$ and stored. Rotation was performed on the magnitude and components of the vector field regions using the function *imrotate* (bilinear interpolation and "crop" mode), implemented in custom codes in Matlab (MathWorks, Inc.). For clarity, the resolution of the vector fields was downscaled using the Matlab function *imresize* with bicubic interpolation.

#### Supplementary Text

#### Supplementary note 1: Active deformation of a bulk nonlinearly elastic solid by a supracellular contractile nematic texture on its free surface

As discussed in the main text, in our system the cellular nematic organization becomes frozen on the free surface of the substrate, both when the adhesive protein is initially homogeneous and isotropic and when it is patterned. For an unpatterned substrate, the process of freezing of the nematic organization has been modeled theoretically by assuming that nematically organized cells deposit nematically aligned matrix, which in turn favors cell alignment (*30*). Here, we focus on situations where the cellular nematic organization has been established, and therefore view it as a nematic texture imprinted onto the free surface of the substrate, very much like the fingerprints in our fingertips. As the solid deforms, the nematic texture deforms with it, consistent with the picture that cell alignment follows adhesive protein alignment, and that these adhesive proteins are attached to the substrate. We assume that the cellular nematic applies active tractions on the substrate as a result of anisotropic contraction along the nematic texture (Fig. SN1).

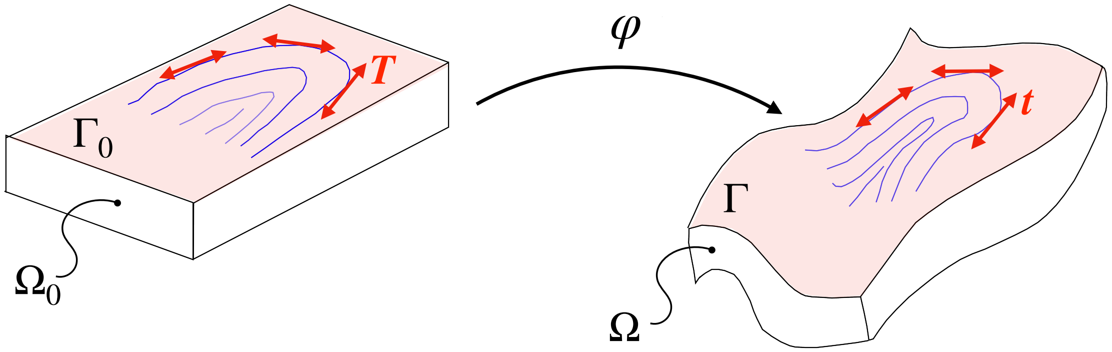


**Figure SN1**: Illustration of the model setup, where a substrate is deformed by the anisotropic contraction of an active nematic texture imprinted on the upper surface.

##### Modeling the bulk elastic substrate

To model this situation mathematically, we consider a 3D body (the substrate) in a reference configuration $\Omega_0$, with part of its boundary $\Gamma_0 \subset \partial\Omega_0$ decorated with the nematic texture. We assume for concreteness that the rest of the boundary is either fixed or traction-free.

We do not restrict the deformations of the substrate to be small, and model it using the theory of nonlinear elasticity (*72, 73*). In a Lagrangian description of the solid substrate, deformation is given by the deformation map $\varphi : \Omega_0 \subset \mathbb{R}^3 \longrightarrow \mathbb{R}^3$. The deformed body and textured surfaces are denoted by $\Omega = \varphi(\Omega_0)$ and $\Gamma = \varphi(\Gamma_0)$. The deformation gradient is the defivative of the deformation map $\boldsymbol{F} = D\varphi$, and its cartesian components are given by ${F^i}_I = \partial\varphi^i/\partial X^I$, with $i = 1, 2, 3$ and $I = 1, 2, 3$. The Cauchy-Green deformation tensor is given by $\boldsymbol{C} = \boldsymbol{F}^T\boldsymbol{F}$ and the Jacobian determinant by $J = \det \boldsymbol{F}$. The elastic energy stored in the substrate can be expressed as

$$\mathcal{F}_{\text{bulk}}[\varphi] = \int_{\Omega_0} W(D\varphi^T D\varphi) dV_0, \tag{S3}$$

where $W(\boldsymbol{C})$ is the stored energy density per unit reference volume or hyperelastic potential. Here, we assume that the substrate is a Neohookean material with

$$W(\boldsymbol{C}) = \frac{\mu_0}{2}\left[\text{tr}(\boldsymbol{C}) - 3\right] - \mu_0 \log(J) + \frac{\lambda_0}{2}\log(J)^2, \tag{S4}$$

where $\mu_0$ and $\lambda_0$ are the Lamé elastic parameters. Using the chain rule, the rate of change of the elastic energy takes the form

$$\dot{\mathcal{F}}_{\text{bulk}}[\dot{\varphi}; \varphi] = \int_{\Omega_0} {P_i}^I \frac{\partial\dot{\varphi}^i}{\partial X^I} dV_0, \tag{S5}$$

where

$$\boldsymbol{P} = 2\boldsymbol{F}\frac{\partial W}{\partial \boldsymbol{C}} \tag{S6}$$

is the first Piola-Kirchhoff stress tensor.

**Modeling the contractile nematic surface texture**

We denote by $\boldsymbol{Q}(\boldsymbol{X})$ for $\boldsymbol{X} \in \Gamma_0$ the traceless and symmetric tensor field characterizing the nematic texture on the surface. In 2D, such a tensor can be expressed as

$$\boldsymbol{Q}(\boldsymbol{X}) = S(\boldsymbol{X})\left[\boldsymbol{T}(\boldsymbol{X}) \otimes \boldsymbol{T}(\boldsymbol{X}) - \frac{1}{2}\boldsymbol{I}\right], \tag{S7}$$

where $S(\boldsymbol{X})$ is the nematic order parameter field and $\boldsymbol{T}(\boldsymbol{X})$ a field of nematic orientations such that $|\boldsymbol{T}| = 1$. We assume for simplicity that active tension in the cellular nematic is uniaxial and contractile along the nematic direction. To model it, let us consider a line of elongated contractile units in the undeformed configuration advected by deformation. We describe this line with a curve $\gamma_0$ on $\Gamma_0$, whose image by $\varphi$ is a curve $\gamma$ on $\Gamma$. At a point $\boldsymbol{X}$ on $\gamma_0$, $\boldsymbol{T}(\boldsymbol{X})$ is a tangent vector to the curve. Because $\gamma = \varphi \circ \gamma_0$, it follows that $\boldsymbol{FT}$ is a vector tangent to $\gamma$ and

$$\boldsymbol{t} = \frac{1}{\lambda_{\boldsymbol{T}}}\boldsymbol{FT} \tag{S8}$$

is a unit vector tangent to $\gamma$, where we have introduced the notation $\lambda_{\boldsymbol{T}} = \sqrt{\boldsymbol{T}\cdot\boldsymbol{C}\cdot\boldsymbol{T}}$ for the stretch ratio along $\boldsymbol{T}$. The rate of change of the line element along this curve is given by (*72, 73*)

$$\frac{Dds}{Dt} = \boldsymbol{t}\cdot\boldsymbol{d}\cdot\boldsymbol{t}\,ds, \tag{S9}$$

where $\boldsymbol{d}$ is the rate-of-deformation tensor at the surface of the body. Then, we can write the power input of the contractile curve $\gamma$ as

$$\mathcal{P}_\gamma = \int_\gamma \tau\omega\,\boldsymbol{t}\cdot\boldsymbol{d}\cdot\boldsymbol{t}\,ds, \tag{S10}$$

where $\tau$ is the line tension per contractile unit and $\omega$ the number of contractile units per unit length along $\gamma$.

Our aim is to express this power input in the reference configuration to have a fully Lagrangian formulation that facilitates the computer implementation of the theory. Accordingly, noting that $\boldsymbol{F}^T\boldsymbol{dF} = (1/2)\dot{\boldsymbol{C}}$ where $\dot{\boldsymbol{C}}$ is the material time derivative the the Cauchy-Green deformation tensor, we express the rate of elongation of the curve as

$$\boldsymbol{t}\cdot\boldsymbol{d}\cdot\boldsymbol{t} = \frac{1}{2}\frac{\boldsymbol{T}\cdot\dot{\boldsymbol{C}}\cdot\boldsymbol{T}}{\boldsymbol{T}\cdot\boldsymbol{C}\cdot\boldsymbol{T}}. \tag{S11}$$

Furthermore, our assumption that contractile units are advected by deformation implies that $\omega ds = \omega_0 ds_0$, where $\omega_0$ is the fixed number of contractile units per unit length along the reference curve $\gamma_0$ and $ds_0$ its line element. By the change of variables formula for integrals, we conclude that

$$\mathcal{P}_\gamma = \int_{\gamma_0} \frac{\tau\omega_0}{2}\frac{\boldsymbol{T}\cdot\dot{\boldsymbol{C}}\cdot\boldsymbol{T}}{\boldsymbol{T}\cdot\boldsymbol{C}\cdot\boldsymbol{T}}\,ds_0 = \int_{\gamma_0} \tau\omega_0\frac{\partial}{\partial t}\left(\log\lambda_{\boldsymbol{T}}\right)\,ds_0. \tag{S12}$$

Assuming that the reference surface is decorated by an ensemble to such contractile lines separated locally by a distance $t_0$, we can express the density of contractile units per unit reference area as $\rho_0 = \omega_0/t_0$. Introducing the uniaxial active tension in the tissue as $\sigma = \tau\rho_0$, we conclude that the power input functional for the ensemble of contractile lines distributed over the surface $\Gamma_0$ can be expressed as

$$\mathcal{P}_{\text{contract}}[\dot{\varphi};\varphi] = \int_{\Gamma_0} \sigma \frac{\partial}{\partial t} \left(\log \lambda_{\boldsymbol{T}}\right) \, dS_0. \tag{S13}$$

This expression shows that, whenever $\sigma$ is constant in time and independent of other variables, the contractile power input can be derived as the rate of change of the anisotropic pseudo-energy functional

$$\mathcal{F}_{\text{contract}}[\varphi] = \int_{\Gamma_0} \sigma \log \lambda_{\boldsymbol{T}} \, dS_0. \tag{S14}$$

##### Governing equations

The governing equations can be obtained by minimizing the Rayleighian functional (*74–76*)

$$\mathcal{R}[\dot{\varphi};\varphi] = \dot{\mathcal{F}}_{\text{bulk}}[\dot{\varphi};\varphi] + \mathcal{P}_{\text{contract}}[\dot{\varphi};\varphi] \tag{S15}$$

with respect to the velocity $\dot{\varphi}$, or equivalently, minimizing $\mathcal{F}_{\text{bulk}}[\varphi] + \mathcal{F}_{\text{contract}}[\varphi]$ with respect to $\varphi$. These minimizations should account for fixed displacement boundary conditions.

Although the Euler-Lagrange equations resulting from these variational statements are not used in the finite element approximation, we provide them for completeness. Integration by parts on the bulk and the surface leads to the statement of balance of linear momentum in the bulk

$$\frac{\partial P_i{}^I}{\partial X^I} = 0 \quad \text{in} \;\; \Omega_0, \tag{S16}$$

and balance of linear momentum at the surface between the solid tractions and those induced by the contractile cellular nematic, which takes the form

$$\boldsymbol{P} \cdot \boldsymbol{N} = \text{Div}_{\Gamma_0} \left( \frac{\sigma}{\lambda_{\boldsymbol{T}}} \boldsymbol{t} \otimes \boldsymbol{T} \right) \quad \text{on} \;\; \Gamma_0. \tag{S17}$$

In this equation, $\boldsymbol{N}$ is the outer normal at the surface, $\text{Div}_{\Gamma_0}$ is the surface divergence, and the term between brackets is the first Piola-Kirchhoff surface tension induced by the active layer.

##### Parameters of the simulations in Fig. 4

We consider a cuboidal geometry for the substrate with dimensions $A \times A \times H$, where $H$ is the thickness. We fix the bottom of the slab with Dirichlet boundary conditions, while the lateral faces are traction-free.

In addition to these dimensions, the model parameters are the Lamé parameters of the hyperelastic model $\lambda_0$ and $\mu_0$, the active tension parameter $\sigma$, the typical radius of $\Gamma_0$, denoted by $R_0$, and the length-scale of nematic defects, $\ell_{\text{defect}}$, introduced later. The resulting dimensionless parameters are $H/R_0$, $A/R_0$, $\ell_{\text{defect}}/R_0$, $\lambda_0/\mu_0$ and $\sigma/(\mu_0 R_0)$. For the simulations in Fig. 4, we estimate from experimental measurements $H/R_0 = 0.1$, $\ell_{\text{defect}}/R_0 = 0.01$, $\sigma/(\mu_0 R_0) = 0.6$. We choose $A/R_0 = 1.5$ to strike a balance between computational cost and independence from boundary conditions, and choose $\lambda_0/\mu_0 = 4.0$ to be reasonably close to an incompressible limit without incurring in volumetric locking.

##### Numerical solution

To numerically approximate this problem, we use trilinear finite elements on a structured mesh of parallepipeds of the substrate. The treatment of $\mathcal{F}_{\text{bulk}}$ is standard. The treatment of the contractile layer is also direct. It is sufficient to we use Gaussian quadrature on the surface mesh of quadrilateral elements of $\Gamma_0$ resulting from the bulk mesh to evaluate $\mathcal{F}_{\text{contract}}$ and its variations. The resulting nonlinear system of equations is solved using Newton's method.

#### Supplementary note 2: Computational generation of nematic patterns

To generate computationally patterns of nematic field with predefined location of defects, we find the ground state of a nematic system with restrains that favor the location of defects. We consider the Landau-de Gennes nematic free energy in the one-constant approximation (*9*)

$$\mathcal{F}_{\mathrm{LdG}}[\boldsymbol{Q}] = \int_{\Gamma_0} \frac{L}{2} |\nabla_0 \boldsymbol{Q}|^2 dS_0 + \int_{\Gamma_0} \phi(\boldsymbol{Q}) dS_0, \tag{S18}$$

where $L$ is the Frank constant, $\phi(\boldsymbol{Q}) = aS^2/2 + bS^4/8$, and $a < 0$ and $b > 0$ are susceptibility coefficients favoring nematic order. The typical length-scale of nematic defects is $\ell_{\mathrm{defect}} = \sqrt{-L/(2a)}$. To favor the location of defects, we introduce the restraint potential

$$\mathcal{F}_{\mathrm{defects}}[\boldsymbol{Q}] = \sum_i \int_{D_i} \frac{K}{2} |\boldsymbol{Q} - \bar{\boldsymbol{Q}}_i|^2 dS_0, \tag{S19}$$

where $i$ labels the different defects, $D_i$ is a circular region of size commensurate to $\ell_{\mathrm{defect}}$, $K$ is a penalty coefficient, and $\bar{\boldsymbol{Q}}_i(\boldsymbol{X})$ is a given field of a typical +1/2 or -1/2 defect of the system. To find the nematic field, we minimize $\mathcal{F}_{\mathrm{LdG}}[\boldsymbol{Q}] + \mathcal{F}_{\mathrm{defects}}[\boldsymbol{Q}]$ with respect to $\boldsymbol{Q}$ with parallel anchoring conditions at the boundary of $\Gamma_0$. Numerically, we solve these equations using linear finite elements. Given $\boldsymbol{Q}$ at each point, we identify $\boldsymbol{T}$ as the eigenvector with positive eigenvalue (Fig. SN2).

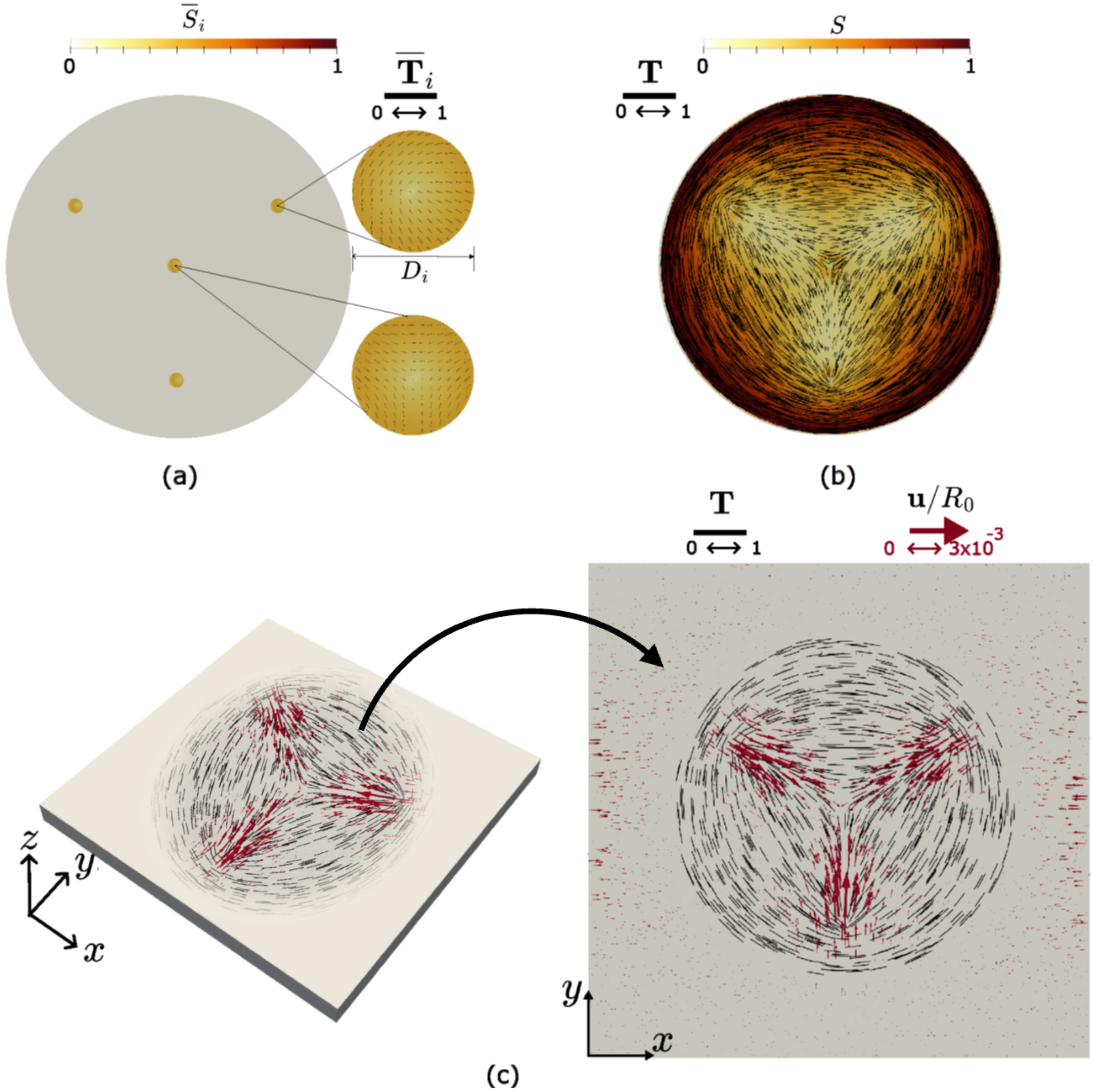


**Figure SN2**: (a) Imposed orientation field in the domain, defined by $\overline{\mathbf{Q}}_i = \overline{S}_i \left( \overline{\mathbf{T}}_i \otimes \overline{\mathbf{T}}_i - \frac{1}{2}\mathcal{I} \right)$. (b) Resulting orientation field obtained by minimizing the Landau–de Gennes nematic free energy $\mathcal{F}_{\mathrm{LdG}}(\mathbf{Q})$; (c) Deformations in the substrate: (left) 3D view of the deformed substrate; (right) top view showing the displacement field $\mathbf{u}/R_0$.

#### Supplementary note 3: Active deformation of a thin shell by a supracellular contractile nematic texture

We consider now the situation where cells are detached from the substrate. Mechanically, we can think of the cell layer as being composed of a passive component, representing for instance ECM remaining beneath of between cells or some cellular organelles, and an active contractile component. Accordingly, we idealize the system as a thin nonlinearly elastic shell with an imprinted contractile nematic texture (Fig. SN3). The latter is modeled as in the previous section.

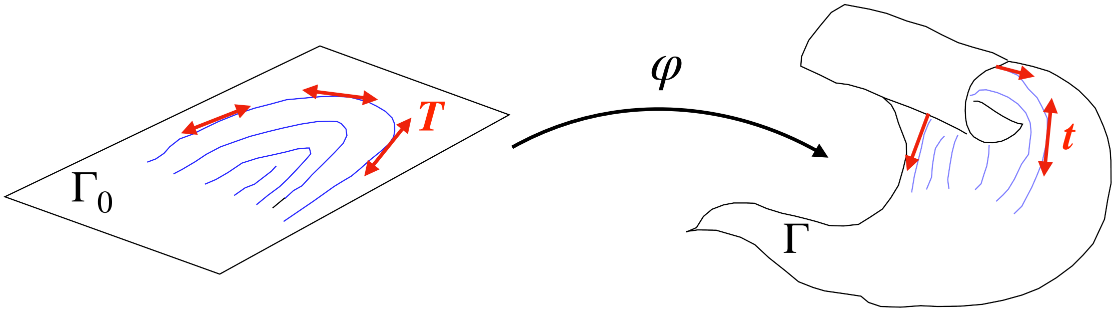


**Figure SN3**: Illustration of the model setup, where a thin shell is deformed by the anisotropic contraction of an active nematic texture imprinted on the thin material sheet.

##### Kinematics

Now, the deformation map of such a shell following detachment is a mapping $\varphi$ from the reference configuration of $\Gamma_0$ into the deformed surface $\Gamma$. For simplicity, we assume that $\Gamma_0$ belongs to the $(X_1, X_2)$ plane described with Cartesian coordinates. We define the natural (or convected) basis of tangent vectors to $\Gamma$ induced by the deformation as

$$\boldsymbol{g}_\alpha = \frac{\partial \varphi}{\partial X^\alpha}, \tag{S20}$$

where greek indices run from 1 to 2. The unit normal field is given by

$$\boldsymbol{n} = \frac{1}{|\boldsymbol{g}_1 \times \boldsymbol{g}_2|} \boldsymbol{g}_1 \times \boldsymbol{g}_2. \tag{S21}$$

We can define the components of the metric tensor in the convected basis as

$$g_{\alpha\beta} = \boldsymbol{g}_\alpha \cdot \boldsymbol{g}_\beta. \tag{S22}$$

The components of the inverse of the metric tensor, $g^{\alpha\beta}$, allowing us to raise indices, are given by the relations

$$g^{\alpha\gamma} g_{\gamma\beta} = \delta^{\alpha}_{\beta}. \tag{S23}$$

The deformation gradient $\boldsymbol{F}$ expressed in Cartesian coordinates is a $3 \times 2$ matrix. We can compute the components of the Cauchy-Green deformation tensor, a $2 \times 2$ matrix, as $\boldsymbol{C} = \boldsymbol{F}^T \boldsymbol{F}$. Because the columns of $\boldsymbol{F}$ are the vectors $\boldsymbol{g}_1$ and $\boldsymbol{g}_2$, we have that $C_{\alpha\beta} = g_{\alpha\beta}$. We define the surface Jacobian as

$$J = \sqrt{\det g_{\alpha\beta}}. \tag{S24}$$

Finally, we define the coefficients of the second fundamental form in the convected coordinates as

$$\kappa_{\alpha\beta} = -\frac{\partial^2 \varphi}{\partial X^\alpha \partial X^\beta} \cdot \boldsymbol{n}. \tag{S25}$$

**Elastic energy of the shell**

We build our model on the standard nonlinear Koiter shell model (*77, 78*). In this model, given the fourth-order isotropic tensor of elastic parameters

$$\mathbb{C}^{\alpha\beta\gamma\delta} = \frac{2\lambda\mu}{\lambda + 2\mu} \delta^{\alpha\beta} \delta^{\gamma\delta} + \mu \left( \delta^{\alpha\gamma} \delta^{\beta\delta} + \delta^{\alpha\delta} \delta^{\beta\gamma} \right). \tag{S26}$$

expressed in terms of the Lamé parameters $\lambda$ and $\mu$, and given the membrane strain

$$\varepsilon_{\alpha\beta} = \frac{1}{2} \left( g_{\alpha\beta} - \delta_{\alpha\beta} \right), \tag{S27}$$

the elastic energy density is given by

$$w(\boldsymbol{\varepsilon}, \boldsymbol{\rho}) = \frac{\mathbb{C}^{\alpha\beta\gamma\delta}}{2} \left( h_0 \varepsilon_{\alpha\beta} \varepsilon_{\gamma\delta} + \frac{h_0^3}{12} \kappa_{\alpha\beta} \kappa_{\gamma\delta} \right), \tag{S28}$$

where $h_0$ is the shell thickness in its reference configuration. The total elastic energy of the shell is then

$$\mathcal{F}_{\text{Koiter}}[\varphi] = \int_{\Gamma_0} w(\boldsymbol{\varepsilon}, \boldsymbol{\rho}) \, dS_0. \tag{S29}$$

The membrane part of this elastic energy is the standard Kirchhoff-Saint Venant hyperelastic model, well-known to produce unphysical material instabilities under compression. Usually, this is not a concern for thin shells, which cannot support significant compression before buckling. However, here the shell is pre-stressed and may undergo significant compression prior to buckling. For this reason, we replace the in-plane part of the Koiter model by a well-behaved 2D Neohookean model given by

$$W(\boldsymbol{C}) = \frac{\lambda^{2D}}{2} (\log J)^2 - \mu^{2D} \log J + \frac{\mu^{2D}}{2} (\operatorname{tr} \boldsymbol{C} - 2) , \tag{S30}$$

where $\lambda^{2D} = h_0\lambda$ and $\mu^{2D} = h_0\mu$ are Lamé parameters integrated through-the-thickness with units of surface tension. The membrane elastic energy is then

$$\mathcal{F}_{\text{membrane}}[\varphi] = \int_{\Gamma_0} W(\boldsymbol{C})\, dS_0. \tag{S31}$$

To define the bending energy, we introduce the fourth-order tensor of elastic moduli as

$$\mathbb{C}^{\alpha\beta\gamma\delta} = \lambda^{2D} \delta^{\alpha\beta}\delta^{\gamma\delta} + \mu^{2D} \left(\delta^{\alpha\gamma}\delta^{\beta\delta} + \delta^{\alpha\delta}\delta^{\beta\gamma}\right), \tag{S32}$$

leading to

$$\mathcal{F}_{\text{bending}}[\varphi] = \int_{\Gamma_0} \frac{h_0^2}{24} \mathbb{C}^{\alpha\beta\gamma\delta} \kappa_{\alpha\beta}\kappa_{\gamma\delta}\, dS_0. \tag{S33}$$

To account phenomenologically for the increase bending stiffness as the tissue contracts and thickens, we can assume cell incompressibility to estimate the current thickness as $h = h_0/J$ and use it instead of $h_0$ in the equation above.

##### Rayleighian functional and dimensionless numbers

To form the Rayleighian functional, we combine the membrane and bending energy functionals with an active power and a dissipation functional. As in the previous section, we define the power input functional $\mathcal{P}_{\text{contract}}[\dot{\varphi}; \varphi]$ as in Eq. (S13). We simplify the dissipative environment of the tissue considering a local viscous drag proportional to the shell velocity. Accordingly, we introduce the dissipation functional

$$\mathcal{D}[\dot{\varphi}; \varphi] = \int_{\Gamma} \frac{\xi}{2} |\dot{\varphi}|^2\, dS = \int_{\Gamma_0} \frac{\xi}{2} |\dot{\varphi}|^2 J\, dS_0, \tag{S34}$$

where $\xi$ is a viscous drag coefficient. With all these ingredients, the dynamics of the system result from minimizing the Rayleighian functional

$$\mathcal{R}[\dot{\varphi}; \varphi] = \dot{\mathcal{F}}_{\text{membrane}}[\dot{\varphi}; \varphi] + \dot{\mathcal{F}}_{\text{bending}}[\dot{\varphi}; \varphi] + \mathcal{P}_{\text{contract}}[\dot{\varphi}; \varphi] + \mathcal{D}[\dot{\varphi}; \varphi] \tag{S35}$$

with respect to $\dot{\varphi}$. Rather than deriving the Euler-Lagrange equations, we outline below how we use this variational formalism to derive the finite element discrete equations.

We are now in a position to enumerate the parameters and key dimensionless numbers of the model. The input parameters for this model are the shape of the domain $\Gamma_0$ with typical radius $R_0$, its nematic texture, the 2D Lame parameters $\lambda^{2D}$ and $\mu^{2D}$, the shell thickness $h_0$, the activity parameter $\sigma$, the drag coefficient $\xi$, and possibly the time-scale of mechanical loading of the system $t_\mathrm{d}$. Physically, this time-scale is related to the detachment rate. Here, we increase active tension from zero to the baseline $\sigma$ using a linear ramp of duration $t_\mathrm{d}$. The dimensionless parameters are the ratio of Lamé parameters $\Pi_1 = \lambda^{2D}/\mu^{2D}$, the elastocapillary number $\Pi_2 = \sigma/\mu^{2D}$, the slenderness $\Pi_3 = 2R_0/h_0$, and the ratio between the loading rate and the bending relaxation time in the dissipative medium $\Pi_4 = t_\mathrm{d}\mu^{2D}h_0^2/(\xi R_0^4)$.

**Table S1**: Model parameters for the study of nematically guided morphogenesis

| | $\lambda^{2D}/\mu^{2D}$ | $\sigma/\mu^{2D}$ | $2R_0/h_0$ | $h_0$ | $\Pi_4$ |
|---|---|---|---|---|---|
| Fig 5. A-top (Movie S8, S10, S14, S16) | 0.5 | 0.9 | 30 | 0.024 | 0.3429 |
| Fig 5. A-bottom (Movie S12) | 0.5 | 0.9 | 60 | 0.024 | 0.0214 |
| Fig 5. D (Movie S18) | 0.5 | 0.9 | 30 | 0.024 | 0.3429 |
| Fig 5. G (Movie S20) | 0.5 | 0.9 | 30 | 0.024 | 0.3429 |
| Fig 6. A (Movie S23) | 0.5 | 0.8 | 30 | 0.024 | 0.3429 |
| Fig 6. B (Movie S25) | 0.5 | 0.7 | 50 | 0.022 | 0.0529 |
| Fig 6. C (Movie S28) | 0.5 | 0.9 | 130 | 0.014 | 0.0029 |
| Fig 7 B (Movie S28) | 0.5 | 0.9 | 130 | 0.014 | 0.0029 |
| Fig 7 C | 0.5 | 0.9 | 130 | 0.014 | 0.0029 |

##### Time-incremental Rayleighian functional

To discretize the problem in time, we consider a sequence of time-instants $t^0, t^1, \ldots, t^n, t^{n+1} \ldots$, and denote the time-step by $\Delta t = t^{n+1} - t^n$, which in principle can be non-constant. We discretize the velocity of the shell as

$$\dot{\varphi}^{n+1} \approx \frac{\varphi^{n+1} - \varphi^n}{\Delta t}. \tag{S36}$$

We approximate the rate of change of the elastic energies as $\dot{\mathcal{F}}^{n+1} \approx (\mathcal{F}^{n+1} - \mathcal{F}^n)/\Delta t$. When solving for $\varphi^{n+1}$, $\mathcal{F}^n$ is constant and can therefore be ignored. Introducing these approximations into Eq. (S35) and multiplying by $\Delta t$, we obtain the incremental Rayleighian as

$$\Delta\mathcal{R}[\varphi^{n+1};\varphi^n] = \mathcal{F}_{\text{membrane}}[\varphi^{n+1}] + \mathcal{F}_{\text{bending}}[\varphi^{n+1}] + \int_{\Gamma_0} \sigma \log \lambda_{\boldsymbol{T}}^{n+1} dS_0 + \int_{\Gamma_0} \frac{\xi}{2\Delta t}|\varphi^{n+1} - \varphi^n|^2 J^n \, dS_0. \tag{S37}$$

Given $\varphi^n$, the time-discrete evolution problem is to find $\varphi^{n+1}$ as the minimizer of $\Delta\mathcal{R}$ with respect to its first argument.

##### Finite element discretization

We approximate in space the variational problem of minimizing $\Delta\mathcal{R}[\varphi^{n+1};\varphi^n]$ with respect to $\varphi^{n+1}$ using surface finite elements to approximate $\varphi^{n+1}(\boldsymbol{X}) \approx \sum_I B_I(\boldsymbol{X})\boldsymbol{x}_I^{n+1}$, where $B_I(\boldsymbol{X})$ are the basis functions supported on a triangulation of $\Gamma_0$, and $\boldsymbol{x}_I^{n+1} \in \mathbb{R}^3$ are the nodal coefficients. Because the bending free-energy depends on the second fundamental form, which in turn depends on second derivatives of $\varphi^{n+1}(\boldsymbol{X})$, the approximation space needs to have square-integrable second derivatives. For that purpose, we use subdivision finite elements based on Loop's scheme (*79*)

##### A simple analytical example

We provide here an idealized application of the theory, amenable to an almost analytical treatment. Consider that $\Gamma_0$ is a planar disk of radius $R_0$ and a family of equispaced and equally contractile concentric circles as shown in the figure. We assume the system remains axisymmetric.

Let us consider a Neohookean in-plane elastic model along with the active contractile lines as described earlier. To simplify the derivation, we model the active contractile lines using a pseudo-elastic energy and we ignore the bending energy. Hence, the total energy of the system is

$$\mathcal{F} = \int_{\Gamma_0} \left\{ \frac{\lambda_0}{2} (\log J)^2 - \mu_0 \log J + \frac{\mu_0}{2} (\operatorname{tr} \boldsymbol{C} - 2) + \sigma \log \lambda_{\boldsymbol{T}} \right\} dS_0. \tag{S38}$$

Because of the symmetry of the problem, the principal stretches of the deformation are along the radial and the angular directions, which we denote by $\lambda_r$ and $\lambda_\theta$. Consequently,

$$J = \lambda_r \lambda_\theta, \quad \operatorname{tr} \boldsymbol{C} = \lambda_r^2 + \lambda_\theta^2. \tag{S39}$$

Furthermore, because of the arrangement of the contractile lines, $\lambda_{\boldsymbol{T}} = \lambda_\theta$. Consequently, the energy density per unit area can be expressed as

$$f = \frac{\lambda_0}{2} (\log \lambda_r + \log \lambda_\theta)^2 - \mu_0 (\log \lambda_r + \log \lambda_\theta) + \frac{\mu_0}{2} \left(\lambda_r^2 + \lambda_\theta^2 - 2\right) + \sigma \log \lambda_\theta. \tag{S40}$$

We now attempt to minimize the free-energy locally, obtaining the equilibrium conditions

$$0 = \frac{\partial f}{\partial \lambda_r} = \lambda_0 (\log \lambda_r + \log \lambda_\theta) \frac{1}{\lambda_r} - \frac{\mu_0}{\lambda_r} + \mu_0 \lambda_r, \tag{S41}$$

and

$$0 = \frac{\partial f}{\partial \lambda_\theta} = \lambda_0 (\log \lambda_r + \log \lambda_\theta) \frac{1}{\lambda_\theta} - \frac{\mu_0}{\lambda_\theta} + \mu_0 \lambda_\theta + \frac{\sigma}{\lambda_\theta}. \tag{S42}$$

Multiplying each of these equations by the corresponding stretches, we obtain

$$0 = \lambda_0 (\log \lambda_r + \log \lambda_\theta) - \mu_0 + \mu_0 \lambda_r^2 \tag{S43}$$

$$0 = \lambda_0 (\log \lambda_r + \log \lambda_\theta) - \mu_0 + \mu_0 \lambda_\theta^2 + \sigma. \tag{S44}$$

Subtracting the first equation from the second, we find that

$$\lambda_\theta^2 - \lambda_r^2 + \frac{\sigma}{\mu_0} = 0, \tag{S45}$$

and hence, requiring the stretches to be positive, we find

$$\lambda_\theta = \sqrt{\lambda_r^2 - \frac{\sigma}{\mu_0}}. \tag{S46}$$

Inserting this equation into Eq. (S44) yields a nonlinear equation for $\lambda_r$. By solving it and evaluating $\lambda_\theta$ from Eq. (S46), we can compute the principal stretches that locally minimize the free energy density given the Lamé parameters ($\lambda_0$, $\mu_0$) and active tension ($\sigma$). These principal stretches define an anisotropic state of "spontaneous in-plane strain" of the elastic sheet in response to the anisotropic tension.

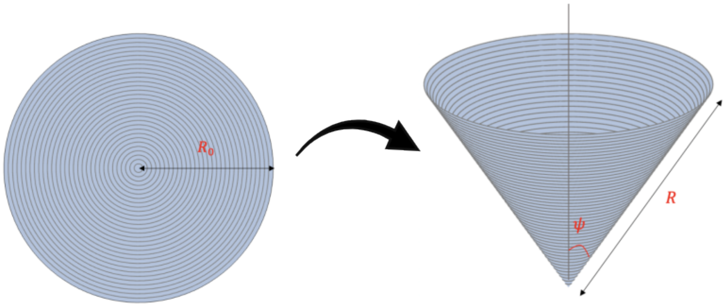


**Figure SN4**: Disk with concentric uniformly spaced contractile lines transforming into a cone.

The question now becomes whether we can find a deformation that realizes these stretches uniformly. The answer is positive and given by a transformation from the disk into a cone with angle $\psi$ and such that a radius of the initial disk is uniformly stretched and becomes a generatrix of the cone with length $R$. Following geometric arguments, we can compute the radial stretch as

$$\lambda_r = \frac{R}{R_0}, \tag{S47}$$

and the angular stretch as

$$\lambda_\theta = \frac{R}{R_0} \sin \psi. \tag{S48}$$

See Fig. SN4 for an illustration. We conclude that given the Lamé parameters ($\lambda_0$, $\mu_0$) and active tension ($\sigma$), it is possible to minimize the energy of the system by deforming the disk into a cone characterized by $R = \lambda_r R_0$ and $\sin \psi = \lambda_\theta / \lambda_r$, where $\lambda_r$ and $\lambda_\theta$ are solutions of Eqs. (S43,S44).

This problem is solved for specific material parameters and the result is plotted on the energy landscape of the system in Fig. SN5. This figure shows that the deformation into a cone with angle $\psi \approx 39°$ has a much lower energy than any flat and isotropic deformation characterized by $\psi = \pi/2$, and hence $\lambda_r = \lambda_\theta$. Consequently, the system will spontaneously buckle out of plane, without any preference for an up- or a down-pointing cone. If bending stiffness is added, then the system will have to overcome a threshold activity for out-of-plane buckling but, if the bending rigidity is small enough, the resulting deformation closely follows the cone geometry with a regularized apex.

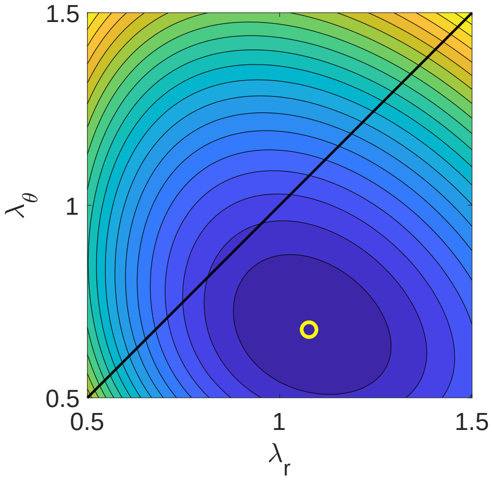


**Figure SN5**: Energy landscape in Eq. (S40) as a function of $\lambda_r$ and $\lambda_\theta$ and equilibrium solution marked by a yellow circle for the parameters $\lambda_0 = 0.5$, $\mu_0 = 1$ and $\gamma = 0.7$. The optimal cone has an angle of $\psi \approx 39°$. The black line represents the $\lambda_r = \lambda_\theta$ corresponding to isotropic and planar deformations, showing that by buckling out-of-plane the system can lower its energy.

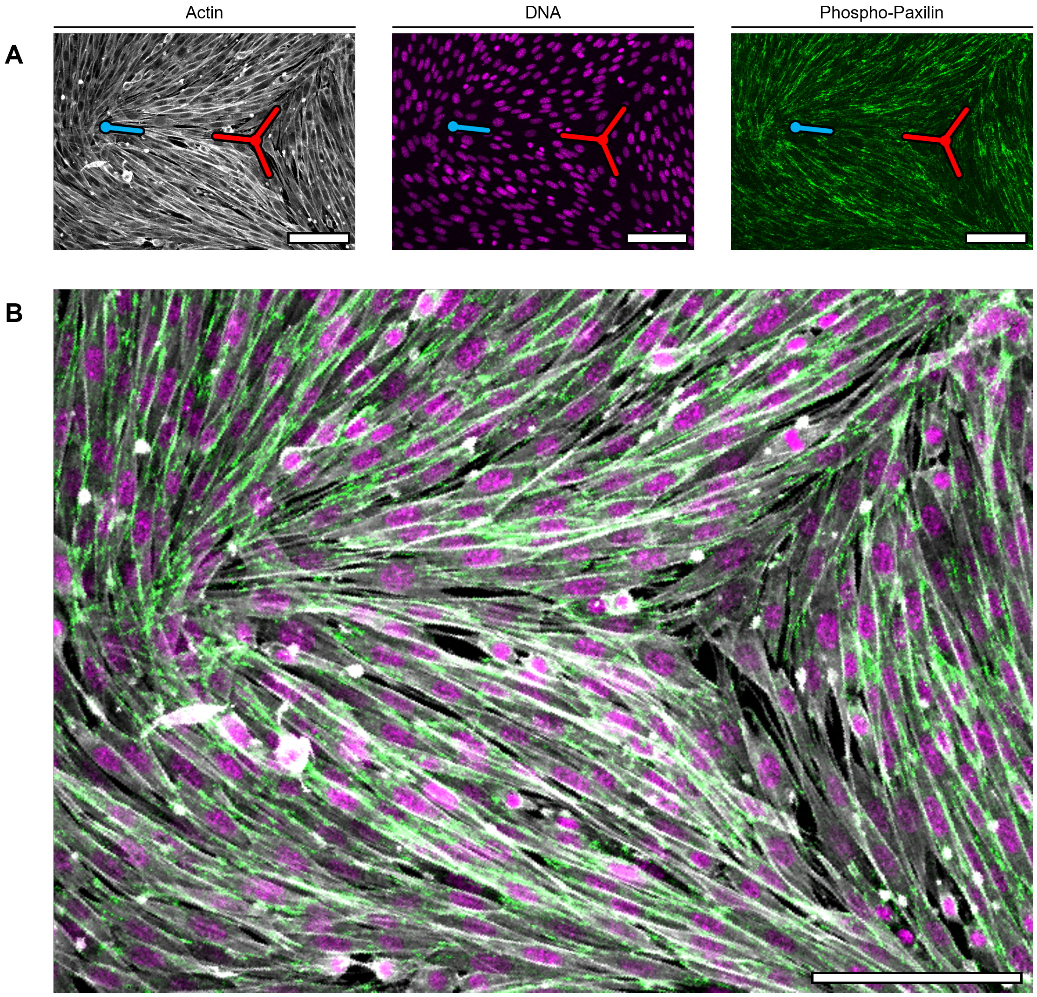


**Fig. S1. Fibroblast nematics.** (**A**) Confocal micrograph of a monolayer of fibroblasts stained with actin, DNA, and phospho-paxilin. Cyan and red symbols indicate the position of positive and negative $\frac{1}{2}$ topological defects, repectively. (**B**) Composite with merged signals from A. Scale bars: 100 $\mu m$.

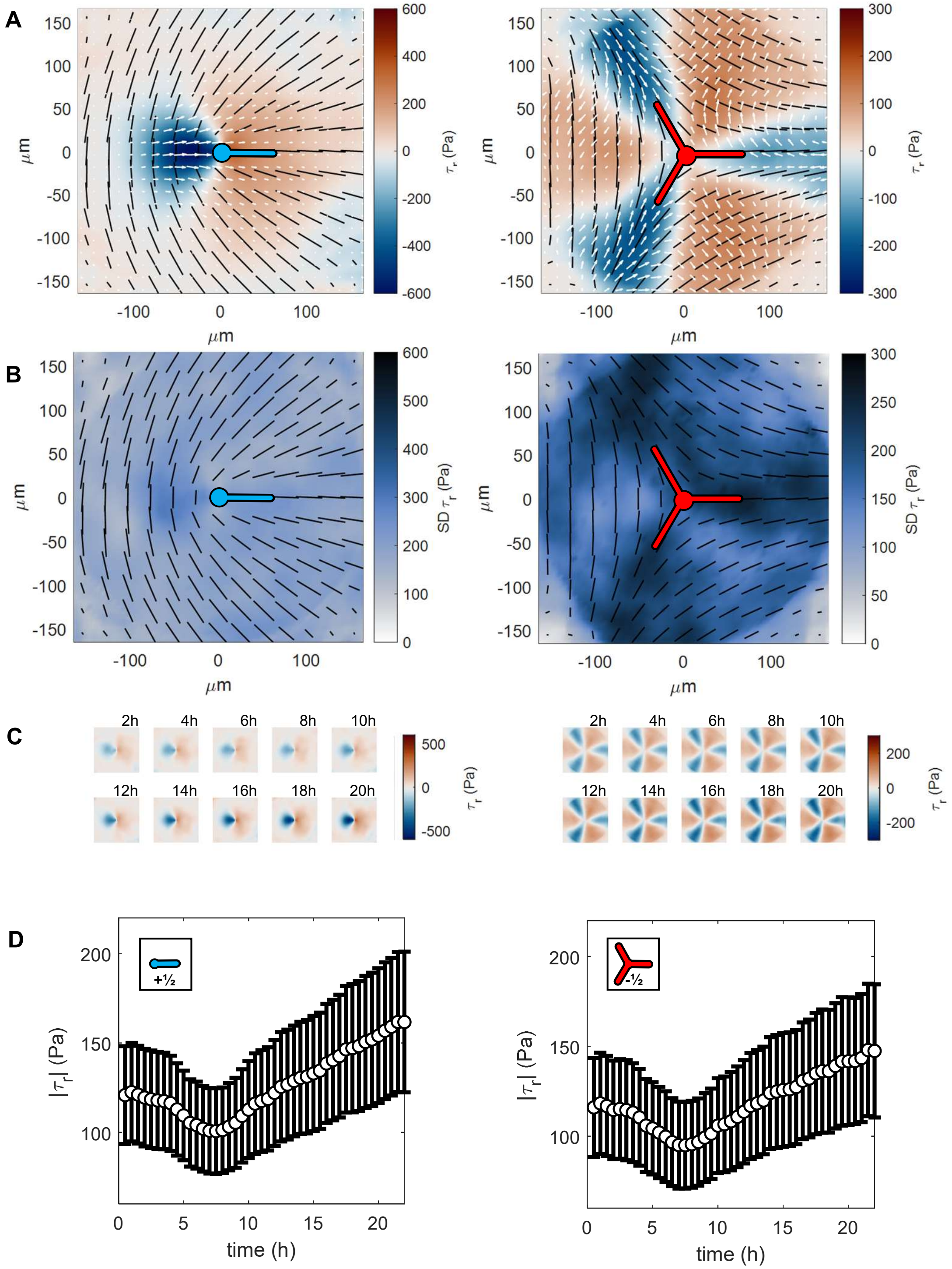

A
B
C
D
μm
τr (Pa)
SD τr (Pa)
2h
4h
6h
8h
10h
12h
14h
16h
18h
20h
|τr| (Pa)
time (h)
+½
-½

**Fig. S2. Defect-mediated traction fields in unconfined fibroblast cellular nematics.** (**A**) Average orientation and traction fields 20h after confluency for $+\frac{1}{2}$ (left) and $-\frac{1}{2}$ (right) topological defects. The black vectors represent the average orientation. The white vectors represent the tractions. The color map shows the magnitude of the radial component of the tractions. (**B**) Standard deviation of the radial component of the tractions in A. (**C**) Color maps showing the time evolution of the radial component of the traction for the $+\frac{1}{2}$ (left) and $-\frac{1}{2}$ (right) topological defects. The initial time point corresponds to the onset of cell confluency in the monolayer. The average fields were obtained with data that spanned a duration of 1 h (N = 15 positions, $n_{+\frac{1}{2}}$=408, $n_{-\frac{1}{2}}$=406 in total). (**D**) Time evolution of the average of the absolute values of the radial component of the traction for $+\frac{1}{2}$ (left) and $-\frac{1}{2}$ (right) topological defects. The error bars correspond to the SEM (N = 15 positions, $n_{+\frac{1}{2}}$=9726, $n_{-\frac{1}{2}}$=9567 in total).

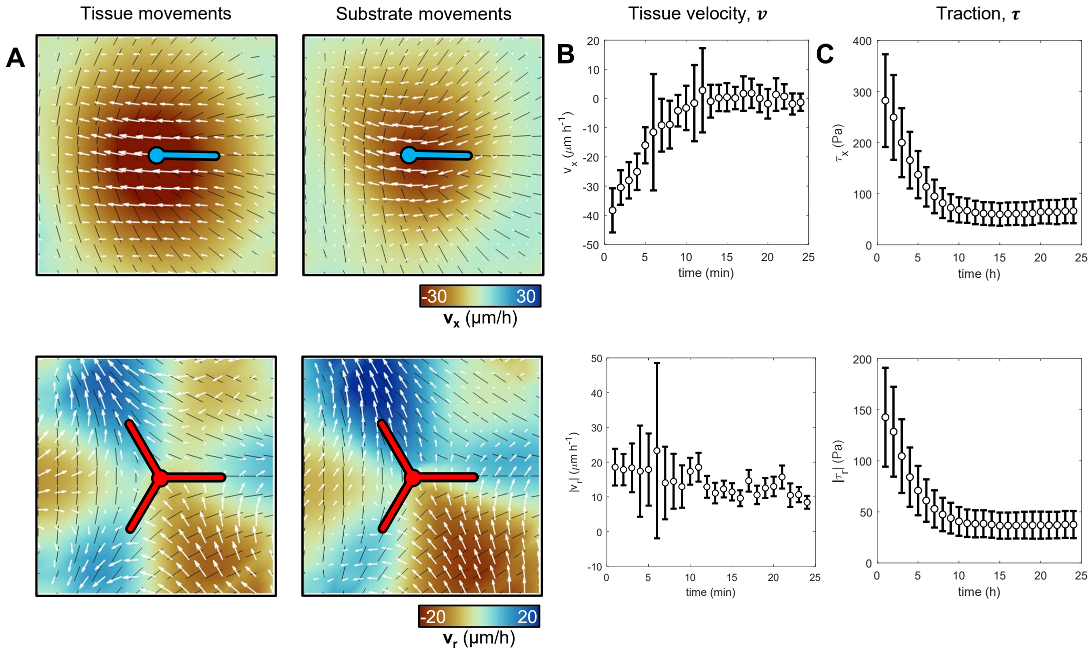


**Fig. S3. Substrate relaxation and defect drag upon inhibition of contractility.** (**A**) Average tissue and substrate movements around $\pm\frac{1}{2}$ topological defects after treatment with Blebbistatin (calculated from 4 timepoints after the treatment corresponding to 2 min). For tissue movements, N=20 positions, $n_{+\frac{1}{2}}$=340, $n_{-\frac{1}{2}}$=215). For substrate movements, N=10 positions, $n_{+\frac{1}{2}}$=115, $n_{-\frac{1}{2}}$=123) Colormaps show the velocity field's horizontal and radial components for positive and negative defects, respectively. (**B**) Time evolution of the horizontal and radial component of the velocity of the tissue in the vicinity of $\pm\frac{1}{2}$ topological defects after the treatment with Blebbistatin ($n_{+\frac{1}{2}}$=340, $n_{-\frac{1}{2}}$=215). (**C**) Time evolution of the horizontal and radial traction components in the vicinity of $\pm\frac{1}{2}$ topological defects after treatment with Blebbistatin. The error bars correspond to the SEM (N=10 positions, $n_{+\frac{1}{2}}$=1032, $n_{-\frac{1}{2}}$=1066).

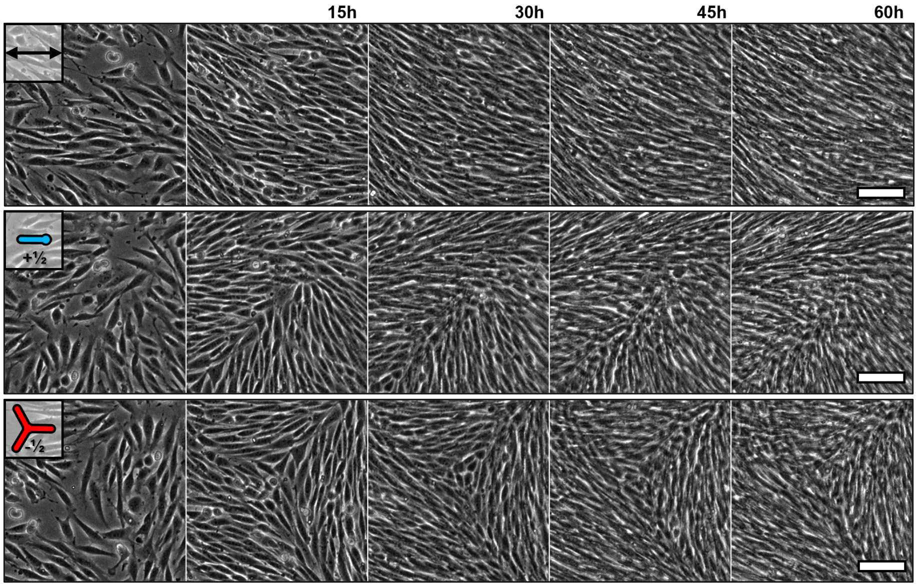


**Fig. S4. Formation of unconstrained nematic fibroblast monolayers** Phase contrast images of regions of an unconfined monolayer of fibroblasts generating, from top to bottom, aligned regions, $+\frac{1}{2}$ and $-\frac{1}{2}$ topological defects. Scale bars: 100 $\mu m$.

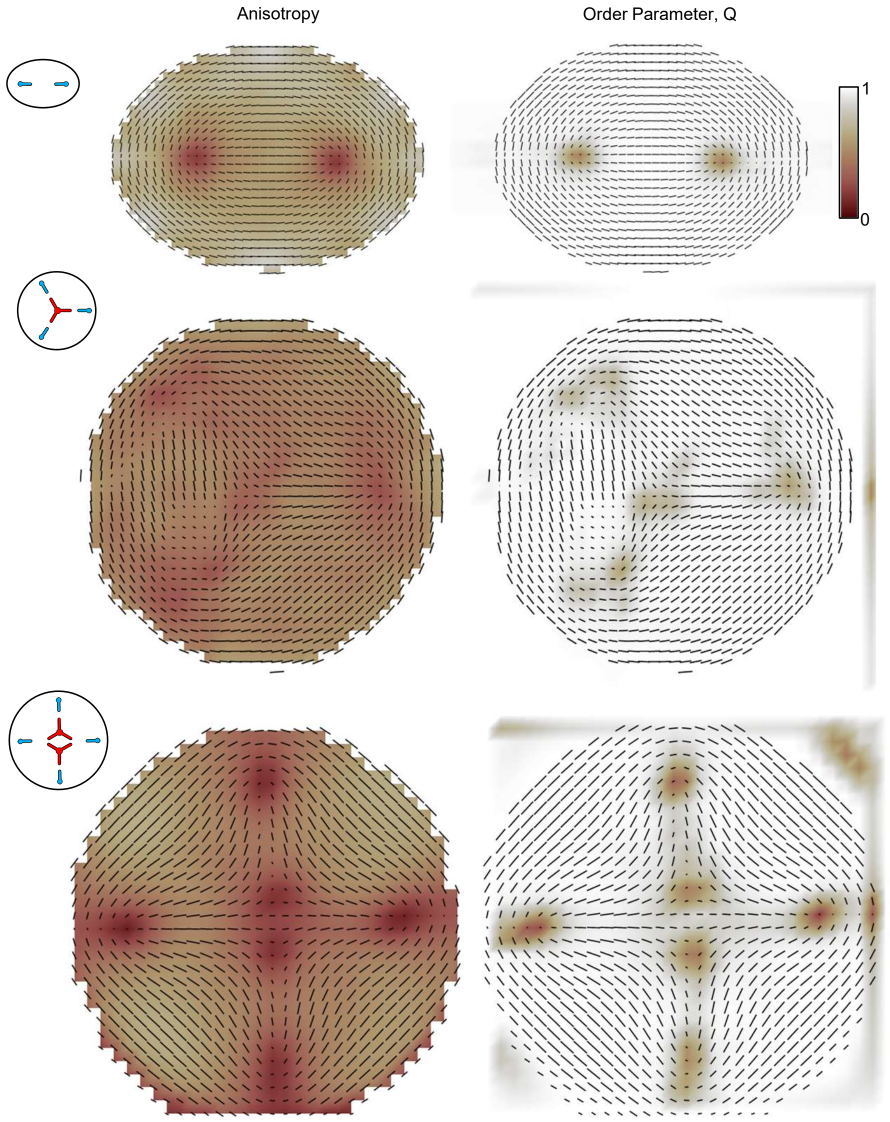
Anisotropy
Order Parameter, Q
1
0

**Fig. S5. Spatial control of nematicity and order parameter.** Average orientation vector field (black vectors) with color maps that represent the average nematicity fields (left) and the order parameter field (right) for 2-, 4-, and 6-topological defect arrangements, shown from top to bottom (n = 29, 5, and 3, respectively).

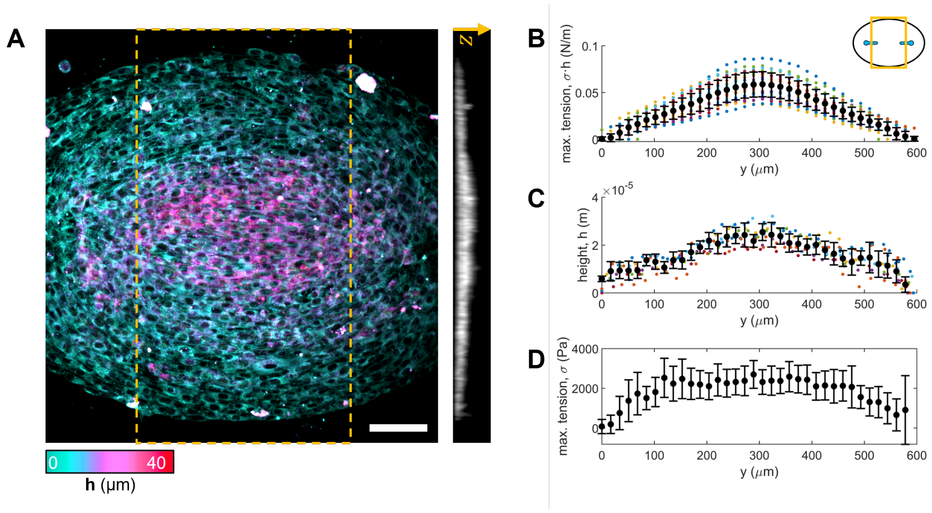


**Fig. S6. Mechanical contribution of cell multilayering in cellular nematics.** (**A**) Maximum Z-projection of a membrane-stained 2-defect cellular nematic. Color indicates the height. Right panel shows the average X-projection of YZ-orthogonal cuts in the regions between the topological defect cores (yellow frame). (**B**) Y-Profile of maximum force per unit length extracted for tension fields of (N=22) 2-defect cellular nematics (inset, Data in Fig. 4C). (**C**) Y-profile of height extracted from the average X-projection of YZ-orthogonal cuts in the regions between the topological defect cores (N=8). (**D**) Y-profile of tension. Scale bars: 100 $\mu m$.

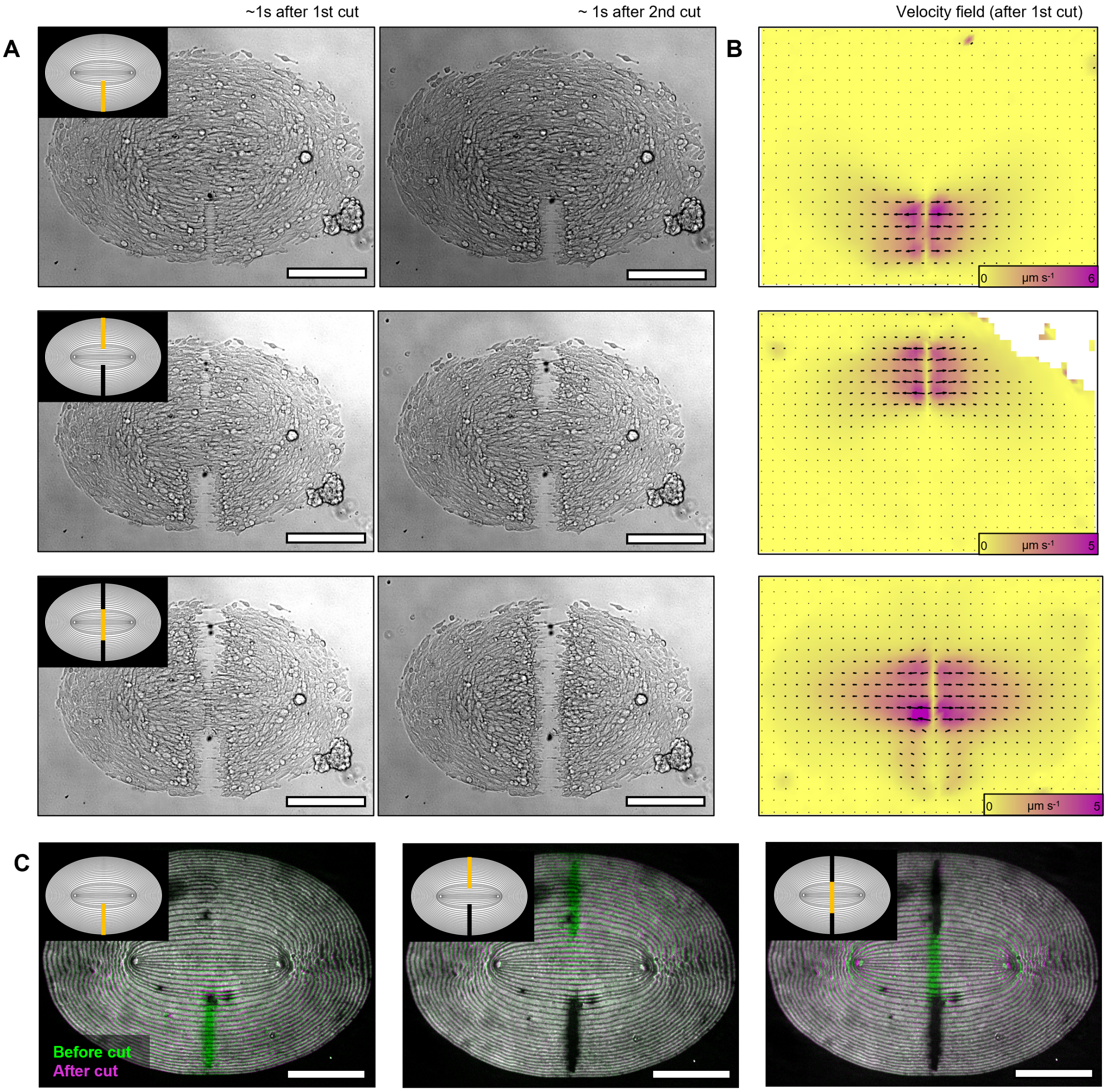


**Fig. S7. Laser ablation of two-defect cellular nematics.** (**A**) Brightfield images of 2-defect cellular nematics 1 second after laser ablation cuts. Inset represents the region of the cut (orange line) and previously cut regions (black lines). (**B**) Velocity field after first cuts in the different positions. (**C**) Fluorescence image composites of the fibronectin micropattern (green) before and (magenta) after laser cuts at different positions. Inset represents the region of the cut (orange line) and previously cut regions (black lines). Scale bars: 200 $\mu m$.

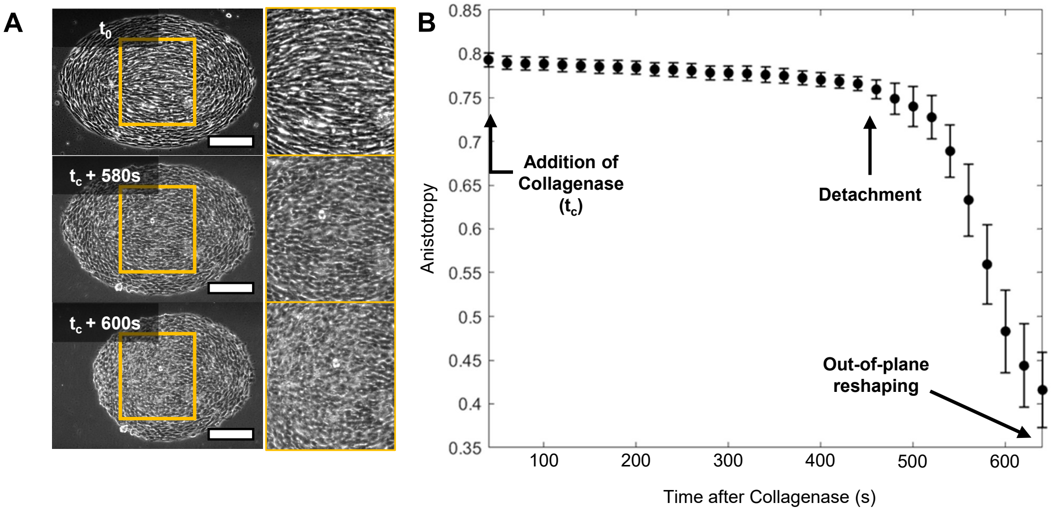


**Fig. S8. Time-evolution of cellular anisotropy during the detachment of minimal cellular nematics.** (**A**) Phase contrast images of 2-defect cellular nematics before ($t_0$) and after the addition of Collagenase, at time $t_c$. The framed regions are magnified for clarity. Framed images correspond to the zoomed region. Scale bars: 100 $\mu m$. (**B**) Variation of normalized average anisotropy in the center of the islands (framed regions) upon the addition of Collagenase. The error bars correspond to the SEM (n=12).

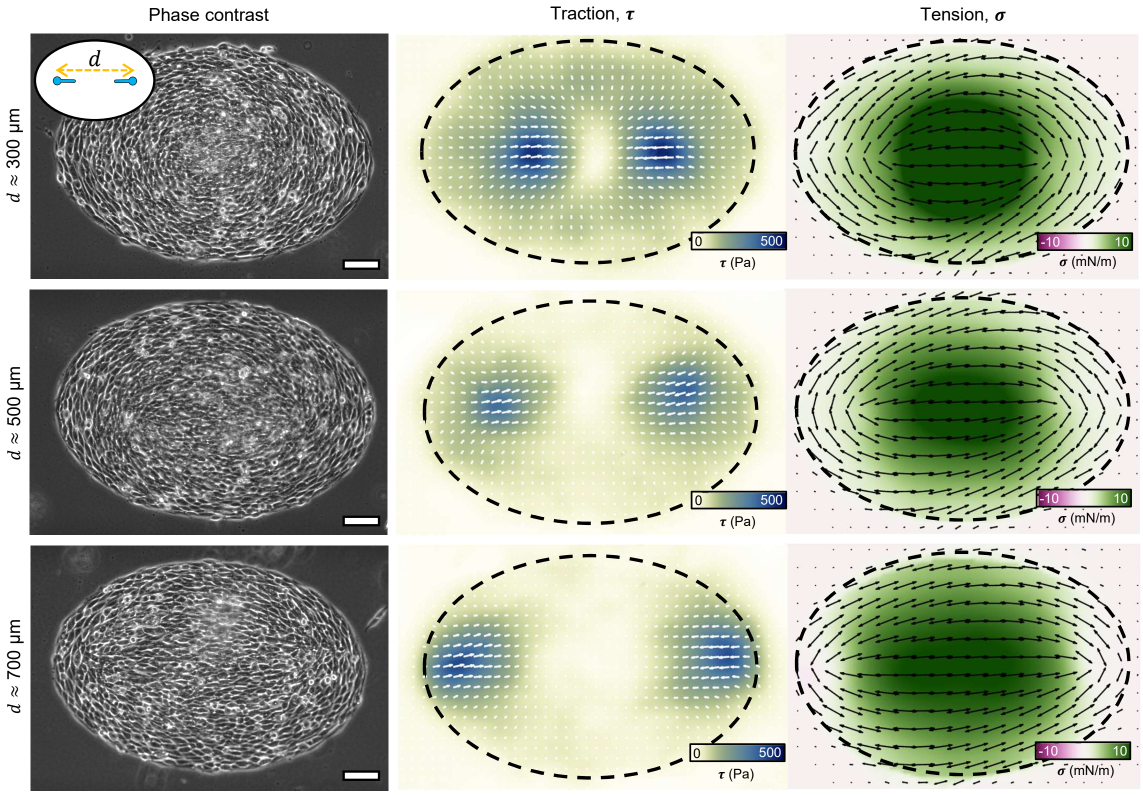


**Fig. S9. Forces by 2-defect cellular nematics featuring different interdefect distances.** Columns, from left to right: Phase contrast image, traction field and tension field. Scale bars: 100 $\mu m$.

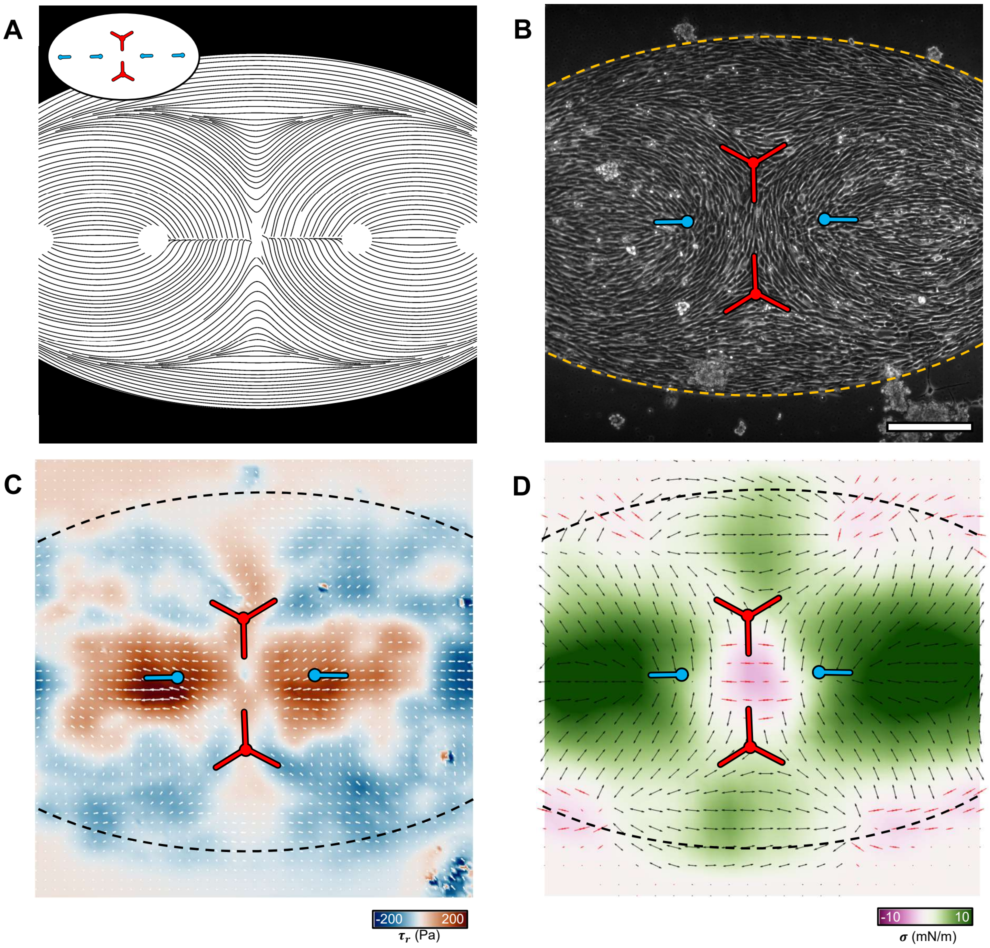


**fig. S10. Compression in 6-defect cellular nematics featuring head-to-head oriented $+\frac{1}{2}$ topological defects.** (**A**) Micropattern design. (**B**) Phase contrast image. Defect positions are indicated with symbols. Dashed line indicate the periphery of the cellular domain. (**C**) Traction field. Colormap indicates the radial tractions with respect to the domain's centroid. (**D**) Tension field. Black arrows indicate tension and red arrows indicate compression. Scale bars: 250 $\mu m$.

**Caption for Movie S1. Phase contrast time-lapse of an unconfined fibroblast nematic on a soft substrate.**

**Caption for Movie S2. Phase contrast time-lapse of an unconfined fibroblast nematic fracturing after Collagenase treatment.** Under high-tension conditions, collagenase treatment resulted in monolayer fracture, typically initiating in regions of low orientational order near -$\frac{1}{2}$ defects.

**Caption for Movie S3. Phase contrast time-lapse of an unconfined fibroblast nematic relaxation after Blebbistatin treatment.** When cellular contractility was reduced using Blebbistatin, the monolayers relaxed, leading to rapid, defect-mediated multicellular motion directed opposite to the defect-characteristic traction fields, driven by the relaxation of the underlying elastic substrate.

**Caption for Movie S4. Phase contrast time-lapse of a Blebbistatin-treated unconfined fibroblast nematic detaching from the substrate after and Collagenase treatment.**

**Caption for Movie S5. Confocal microscopy slice of a wrinkled fibroblast monolayer during multiple laser ablations.**

**Caption for Movie S6. Phase contrast time-lapse of the formation of fibroblast nematics on micropatterned substrates.** From top to bottom, micropatterns impose horizontally aligned orientations, a $+\frac{1}{2}$ topological defect arrangement, and a $-\frac{1}{2}$ topological defect arrangement.

**Caption for Movie S7. Bright-field (left) and epifluorescence (right) time-lapse during laser ablation of 2-defect cellular nematics.**

**Caption for Movie S8. Simulations. Deformation of 2-defect cellular nematics.** The tension magnitude field is homogeneous.

**Caption for Movie S9. Phase contrast time-lapse of the detachment of 2-defect cellular nematics leading to a bowl-shaped structure.** The nuclei (green) at $z = 0\mu m$ are imaged by spinning disk confocal microscopy.

**Caption for Movie S10. Simulations. Comparison of the deformation dynamics of two-defect cellular nematics with homogeneous (left) versus heterogeneous (right) thickness distributions.** In these calculations, active tension is assumed to be proportional to thickness as suggested by the data in Fig. S6.

**Caption for Movie S11. Phase contrast time-lapse of the detachment of a large 2-defect cellular nematics leading to a bowl-shaped structure.**

**Caption for Movie S12. Simulations. Deformation of a large 2-defect cellular nematics.**

**Caption for Movie S13. Phase contrast time-lapse of the detachment of 2-defect cellular nematics with short interdefect distance.**

**Caption for Movie S14. Simulations. Deformation of 2-defect cellular nematics with short interdefect distance.**

**Caption for Movie S15. Phase contrast time-lapse of the detachment of 2-defect cellular nematics with large interdefect distance.**

**Caption for Movie S16. Simulations. Deformation of 2-defect cellular nematics with large interdefect distance.**

**Caption for Movie S17. Phase contrast time-lapse of the detachment of 2-defect cellular nematics with defects rotated 30º.**

**Caption for Movie S18. Simulations. Deformation of 2-defect cellular nematics with defects rotated 30º.**

**Caption for Movie S19. Phase contrast time-lapse of the detachment of 2-defect cellular nematics with defects rotated 90º.**

**Caption for Movie S20. Simulations. Deformation of 2-defect cellular nematics with defects rotated 90º.**

**Caption for Movie S21. Phase contrast time-lapse of the detachment of cellular nematics with spontaneously generated random defect arrangements.**

**Caption for Movie S22. Phase contrast time-lapse of the detachment of 4-defect cellular nematics.**

**Caption for Movie S23. Simulations. Deformation of 4-defect cellular nematics.**

**Caption for Movie S24. Phase contrast time-lapse of the detachment of 6- defect cellular nematics.**

**Caption for Movie S25. Simulations. Deformation of 6-defect cellular nematics.**

**Caption for Movie S26. Phase contrast time-lapse of the detachment of 6- defect cellular nematics with head-to-head oriented $+\frac{1}{2}$ topological defects.**

**Caption for Movie S27. 360º rotation of a 3D-rendered tissue structure.** This shape resulted from the deformation of a 6-defect cellular nematic arrangement, featuring head-to-head oriented $+\frac{1}{2}$ topological defects.

**Caption for Movie S28. Simulations. Deformation of 6-defect cellular nematics with head-to-head oriented $+\frac{1}{2}$ topological defects.**
